# The missing link between biomolecular condensates and amyloid fibrils

**DOI:** 10.64898/2026.01.27.702026

**Authors:** Art Hoti, Shuya Zang, Ephraim Prantl, Michele Grünewald, Alexander Kros, H. Jelger Risselada, G.J. Agur Sevink

## Abstract

The traditional view of protein self-assembly posits a binary choice between phase separation into fluid condensates and nucleation into crystalline amyloid fibers. However, this framework is incomplete. Experiments show that liquid condensates are non-equilibrium systems that mature into solid-like structures mediated by amphiphilic prion-like domains (PLDs). Being spatially organized yet dynamic, lyotropic phases represent an intermediate regime between these states. Using physics-based *de novo* protein design (Evo-MD), we identify a vast amphiphilic motif space encoding fluid lyotropic phases (e.g., micelles and bicelles). TEM, CD, and AlphaFold predictions confirm that these motifs also assemble into amyloid-based hydrogels as thermodynamic endpoints. Notably, the molecular grammar of lyotropic motifs overlaps strongly with that of PLDs and LARKS. Thus, while PLDs likely evolved to stabilize condensates through transient interactions near criticality, our results show that these same amphiphilic forces inherently encode lyotropic structuring and subsequent amyloid formation – linking functional condensation with pathological aggregation.

## 1 Introduction

The ability to understand the origin and role of biomolecular spatio-temporal organization is paramount to our understanding of living systems. Canonical molecular biology posits that lipids are the key molecules that enable this control. Driven by predominantly hydrophobic forces, lipids undergo self-assembly, giving rise to mesoscale, membrane-bound compartments that establish and regulate differences between their internal and external environments. This fundamental role of lipid membranes in cellular organization has long dominated our understanding of biological compartmentalization.

Recent years have witnessed a paradigm shift in our understanding of biological self-organization through the discovery of multiple protein-based assembly mechanisms. Biomolecular condensates have emerged as dynamic, membrane-less organelles formed through the liquid-liquid phase separation of intrinsically disordered proteins (IDPs) [1]. Simultaneously, the discovery of functional amyloids has demonstrated how ordered protein assemblies can serve essential biological roles, challenging traditional views of protein aggregation as purely pathological [2–6].

At first glance, these discoveries suggest a binary landscape of IDP self-assembly. On one hand, disordered assemblies driven by isotropic, multivalent interactions generate homogeneous, liquid-like spherical droplets through liquid-liquid phase separation. On the other, amyloid-prone proteins undergo ordered, nucleation-driven assembly to form highly stable and directional structures stabilized by hydrogen bonding networks. Nevertheless, recent advances in both fields reveal a far more nuanced reality. In particular, the mesoscale architectures of condensates – initially treated by the field as monophasic entities – are now recognised to exhibit tremendous diversity. Computational studies using minimal models of intrinsically disordered proteins have demonstrated that protein sequence patterning alone can encode the formation of an array of structures including membranes, micelles, and worm-like assemblies [7]. From the experimental perspective, multiple structures have been verified, with examples including core-shell configurations [8–10], hierarchical multi-phasic structures [11, 12], micelles[13–15] and even the formation of protein-RNA vesicles with lipid-like properties - including local ordering, size-dependent permeability, and selective encapsulation [16, 17].

From the standpoint of fibril forming motifs, the discovery of Low Complexity Aromatic-Rich Kinked Segments (LARKS) [18] in proteins such as FUS [19, 20], TDP43 [21], and hnRNPA2 [22], further challenges this binary paradigm, demonstrating the existence of assemblies that bridge ordered and disordered states. While sharing structural characteristics with pathogenic amyloids, including cross-*β* motifs stabilized by *π*-*π* stacking and hydrogen bonding, LARKS-based assemblies exhibit distinct properties. Their molecular architecture, featuring proline and glycine induced kinks that prevent tight amyloid-like packing, results in inter-sheet interaction energies of 15-45*k*_B_*T* [12], as opposed to the 50-80 *k*_B_*T* present in pathological fibers. This allows them to form dynamic hydrogel networks that maintain functionality while avoiding irreversible aggregation [20]. Significantly, LARKS are enriched in the low complexity domains of condensate-forming proteins [23] and frequently play a role in condensate maturation from liquid to solid states [12, 24]. Recent studies have revealed that LARKS-containing proteins preferentially assemble at condensate interfaces, forming a mechanically rigid “crust” around a liquid core [25–27]. While this surface-induced assembly has been attributed to enhanced density fluctuations and increased propensity for perpendicular protein alignment at the condensate interface [28], the molecular mechanism underlying the liquid-to-solid transition in protein systems remains poorly understood, with only a few models recently proposed [12, 25, 26, 29]. This phenomenon is a reflection of a more general series of discoveries, which increasingly highlight that the interface between the ‘dilute’ and ‘dense’ phases serves as a crucial site for both functional and pathological control of condensate properties. Pathologically, idemic protein aggregation mechanisms that occur via liquid-liquid phase separation (LLPS), and also the aggregation of condensate host proteins such as *α*-synuclein are catalysed by the condensate surface, leading to aberrant disease-associated states [30]. Functionally, molecular adsorption of amphiphilic proteins to the interface is proposed as a mechanism by which cells can tune and regulate the size of condensates [31, 32]. Additionally, aggregation of the nuclear pore complex (NPC) associated protein Nsp1 at the surface of FG-nup condensate hydrogels has been shown to be critical for the maintenance of their liquid rheology[33, 34]. This complexity in protein assembly is further highlighted by the current state of amyloid research, where the broad diversity of amyloid-like structures, their varying physical properties, and their different propensities to form higher-order structures have led to an ontological crisis in the field [35, 36]. The challenge of categorizing these diverse assemblies underscores the need for a more nuanced framework for understanding protein self-organization.

The continuous transition from liquid condensates to ordered assemblies, particularly evident in LARKS-containing systems, combined with the diverse landscape of protein assemblies observed in nature, raises a fundamental question: what other (intermediate) phases might exist between purely disordered liquid condensates, and highly structured rigid crystalline structures? The recurring observation of protein assemblies that combine order with dynamics suggests the existence of overlooked intermediate states. Such assemblies, which maintain orientational order while preserving dynamic properties, may represent a distinct interconnecting region in the protein phase diagram. Here, we show that intrinsically disordered motifs – whose sequence space strongly overlaps with prion-like sticker motifs in condensate-forming proteins (e.g., FUS, hnRNPA1, hnRNPA2, and TDP43) – can access an overlooked intermediate: lyotropic liquid-crystalline phases. We hypothesize that prion-like motifs, including LARKS, evolved to form transient hydrophobic sticker–sticker interactions that stabilize biomolecular condensates while maintaining overall solubility. This balance is encoded in the amphiphilic sequence of the motif, which integrates attractive hydrophobic and solvating hydrophilic interactions. However, the directed hydrophobic forces required for condensate formation can over-shoot, driving the formation of larger, thermodynamically stable lyotropic assemblies that, under prolonged aging or stress, seed amyloid fibrils.

To systematically test this hypothesis, we employed a genetic algorithm integrated with molecular dynamics simulations [37, 38] to perform a bottom-up, *de novo* exploration of the intrinsically disordered protein (IDP) sequence space encoding lyotropic liquid-crystalline phases. Inspired by amphiphilic lipid assemblies, we optimized sequences for the formation of two-dimensional lamellae, the most generic structural element of lyotropic phases.

## 2 Results

The requirement for balanced amphiphilicity in condensate prion-like domain stickers naturally raises the question of whether such intrinsically disordered motifs can also access more structured, lyotropic phases analogous to those formed by conventional amphiphiles. In this section, we detail results from our evolutionary strategy aimed at the bottom-up *de novo* design of lyotropic phase forming motifs. Our genetic algorithm–guided exploration of lyotropic sequence space is followed by an analysis of the influence of solvent conditions on the resulting phase morphologies. Finally, we demonstrate a strong connection between the sequence-level features and structural organization of lyotropic and prion-like domains [39], using AlphaFold 3 predictions and experimental data to provide insight into the relationship between lyotropic and amyloid phases.

### 2.1 A fitness function for lyotropic phase optimization

We adopt a physics based, inverse-design methodology, using our in-house software Evo-MD [37, 38], that is based on Martini 3 coarse grained molecular dynamics simulations [40]. The utilization of a genetic algorithm enables us to explore the sequence-function space in a *de novo* manner for lyotropic phase forming capacity. The procedure, outlined in Figure 1A and detailed in the methods section consists of the following discrete steps: 1.) Random initialization of the sequence pool (512 sequences), 2.) MD simulations for each sequence, 3.) Evaluation of the fitness function, 4.) Genetic operations cross-over, recombination and random point mutations performed on high-scoring sequences, forming the next generation of peptides. This process is repeated iteratively until fitness convergence is obtained.

**Figure 1.**
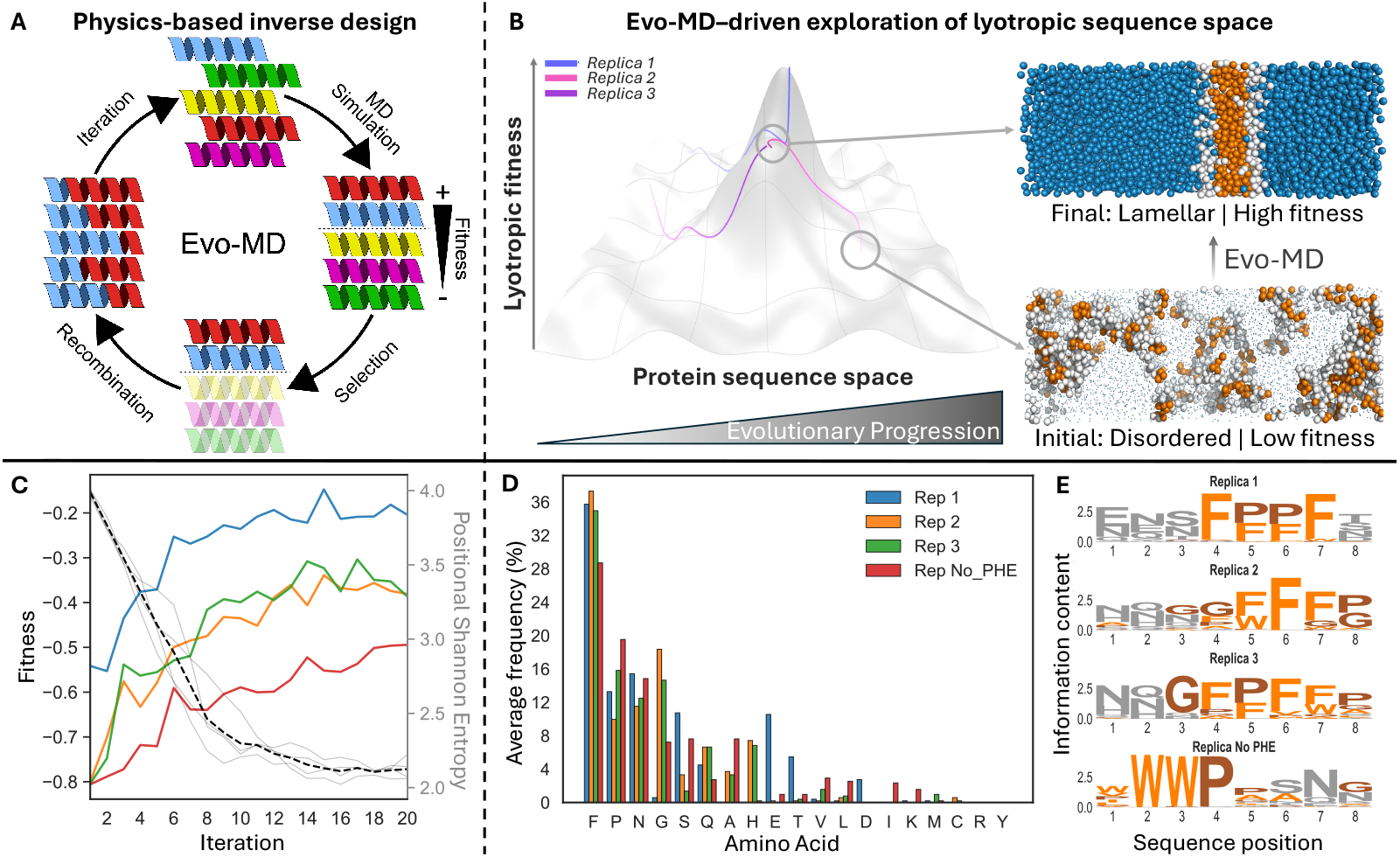
Discovery of the molecular grammar of lyotropic phase-forming sequences. (A) Schematic of the EvoMD workflow, which involves iterative sequence generation, molecular dynamics simulation, fitness evaluation, and genetic operations that guide traversal of sequence space. The *α*-helical peptide representations are purely illustrative, as the sequences of interest are intrinsically disordered. (B) (Left) Conceptual illustration of the lyotropic fitness landscape. Colored trajectories represent independent evolutionary replicas, each initiated from different regions of sequence space (low opacity) and progressing toward optima which may be local or global. (Right) Bead representations showing the evolution of an example disordered phase from the first iteration to an ordered lyotropic lamellar phase from the final iteration. Hydrophobic residues are shown in orange; hydrophilic residues in white. (C) Convergence of four independent EvoMD replicas initiated from different random sequence pools, each containing 512 peptides. Solid colored lines show the fitness of the elite sequence within the iteration; thin gray lines indicate genetic diversity (mean positional Shannon entropy) for each replica, and the dashed line shows the average across replicas. Blue, orange, and green correspond to replicas 1, 2, and 3, respectively. The red line denotes the `No PHE` run, which excludes phenylalanine from the design alphabet. (D) Amino-acid frequency distributions for the top 64 sequences from each replica reveal a common compositional grammar underlying lyotropic phase formation. Note that “F” denotes both F and W aromatic residues. Their relative contributions are distinguished in panel E. (E) Sequence logos of the top 64 peptides per run. Hydrophilic (grey), hydrophobic (orange), and glycine/proline (brown) residues are colored as indicated; logo heights denote positional information content (bits).

The lyotropic-phase fitness function, which determines the sequence space domain that is sampled, is informed by lyotropic phase theory, which holds that if a sequence can form any one of the relevant morphologies – in our case lamellar (L), worm-like micelle (C), or spherical micelle (S) phases (Fig. 3A) – it should, under the appropriate environmental conditions, be capable of transitioning among the others. This principle simplifies the design challenge: rather than targeting all possible lyotropic morphologies simultaneously, we need only identify a single, well-defined point within the lyotropic phase diagram. Lamellar structures exhibit scale-invariant morphology in a slab geometry, and hence provide a clear and unambiguous target for genetic-algorithm optimization independent of system size. As such, we targeted the lamellar phase. Within the slab simulation geometry, the lamellar phase exhibits a clear separation between its hydrophobic and hydrophilic components along the long-axis (which we here select as the z-axis, *L*_*Z*_ *>> L*_*X*_, *L*_*Y*_) (Fig. 1B, top right). This contrasts with a disordered phase scenario where the densities of hydrophobic and hydrophilic amino acids significantly overlap (Fig. 1B, bottom right). As such, the lyotropic phase fitness function minimizes the spatial overlap between hydrophobic and hydrophilic amino acid residues within the system along the long axis (See Methods). The value of the fitness function ranges between −1 and 0, where −1 corresponds to perfect overlap between hydrophobic and hydrophillic residues, i.e. a disordered phase, and 0 corresponds to no overlap between hydrophobic and hydrophillic residues, i.e. a lamellar phase.

The question then becomes: What is the molecular grammar associated to lyotropic phase behaviour?

### 2.2 Inverse design identifies a molecular grammar optimal for lyotropic phase formation

Before analyzing the lyotropic sequence grammar, we first outline two considerations that further narrow down our design objectives. A central question motivating this work is whether the sequence features that promote lyotropic phase formation overlap with those already associated with amyloid formation and biomolecular condensation. The appearance of such an overlap would suggest that the molecular grammar of amyloid- and condensate-forming domains (e.g., prion-like domains [41], LARKS[18]) is shared with lyotropic phase–forming domains. This would imply that proteins previously thought to form only amyloids or condensates may also access lyotropic states under appropriate conditions. Upon self-assembly, the designed motif must satisfy an architectural requirement we refer to as the *loop criterion*, which states that the motif must be short enough to embed as a ‘sticker’ within an intrinsically disordered protein region or loop architecture, and that its termini must reside on the same side of the membrane or interface upon assembly. For this reason, they must evolve to form a multiblock architecture rather than a simple diblock, where one terminus would always be hydrophobic and the other hydrophilic. To ensure the multiblock architecture, we impose a symmetry within the sequence. Particularly, we construct peptides by repeating sequence motifs of length *N*_*m*_, *N/N*_*m*_ times, where *N* is the total number of residues in the chain. To remain within the biologically relevant length range of LARKS-like and amyloid-forming fragments, and to maintain a sufficiently rich solution space 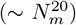, we select a repeat motif length of *N*_*m*_ = 8 and repeat it three times to form a 24 residue peptide. This choice first enables agreement with the *loop criterion* through the formation of multiblock sequences, and second, ensures a length-scale that aligns well with that of established amyloid and “amyloid-like” motifs, thereby enabling direct comparison.

The other consideration is the need for robustness, which we address by performing multiple genetic algorithm runs, referred to as *replicas*. Even when hyperparameters are fixed, each GA run contains several sources of aleatoric uncertainty: the randomly generated initial sequence population, the inherent randomness in mutation, crossover, and selection operations, and the chaotic and non-ergodic nature of the molecular dynamics simulations which are used to evaluate the fitness function [42]. As a result, different runs explore evolutionary trajectories on different parts of the fitness surface, introducing the possibility of convergence to separate local optima. By comparing sequence features that emerge across independent replicas, we additionally obtain information on uniqueness, i.e. whether lyotropic phase formation is governed by a single, well defined molecular grammar or if instead, it can be encoded by multiple distinct grammars that all give rise to the same macroscopic phase behavior. We use the term molecular grammar to denote a broad high-fitness domain in the landscape, within which multiple local optima may exist [43].

Examining EvoMD results, we find that after 20 iterations of optimization (Fig. 1C), replica 1 converged to an optimum with a fitness of −0.20 for the sequence [ENDFFPFT]_3_. On the other hand, replicas 2 and 3 converged to distinct local optima with reduced fitness’s of −0.38 for [NHQGFFFG]_3_ and −0.39 for [HQGFPFWA]_3_, respectively. Because phenylalanine emerges as the most dominant amino acid in the optimal solution space, we also assessed how evolutionary trajectories adapt in its absence by running a replica termed `No PHE`, where phenylalanine was excluded from the alphabet. In `No PHE`, the highest scoring sequence in iteration 20 was [WWWPANNN]_3_, with a fitness of −0.49.

Inspection of the final amino acid frequency distributions (Fig. 1D) and sequence logos (Fig. 1E) for each replica reveals the sequence features required for lyotropic phase formation. Phenylalanine dominates across replicas 1-3 (34.18 ± 3.77 %). On the contrary, in the `No PHE` replica, tryptophan assumes this role, confirming an aromatic hierarchy. Proline (14.65 ± 4.04%) and glycine (10.21 ± 7.91%) form the next group, followed by polar amino acids: arginine (13.6 ± 1.86%), serine (5.76 ± 4.22%), and glutamine (5.13 ± 1.89%). As such, the algorithm consistently selects for a specific functional grammar: aromatics drive aggregation, P/G offer structural kinks/flexibility, and polar residues provide solubility [18]. Hence, all replicas converged to the same grammatical framework, confirming that we have identified the core design principles rather than replica-specific optima. This conclusion is further supported by the observation that pairwise correlations between the amino acid distributions of separate replicas increased over the course of evolution and, by the final iteration, exceeded those between each replica and its respective random initial pool (Fig. S1).

Sequence logos from the final iteration (Fig. 1E) reveal how these building blocks organize into higher-order patterns. Across all replicas, aromatic residues cluster together and are frequently flanked by proline or glycine. Aromatic positions are surrounded by regions of lower information content enriched in polar (hydrophilic) amino acids, suggesting that positions dominated by polar residues tolerate greater sequence variation, whereas aromatic positions require specific amino acid identities. To quantify this, we follow condensate nomenclature and classify aromatic amino acids as “stickers” and hydrophilic residues as “spacers” (Fig. S2). The information content of positions dominated by sticker residues averages 3.584 ± 0.737 bits, whereas positions characterized by hydrophilic spacer residues average 2.452 ± 0.314 bits. This difference is statistically significant (Mann–Whitney U test, *p* = 0.00109). Beyond amino acid composition, we also examined several higher-order physicochemical sequence features. These included sequence charge decoration (SCD [44]), sequence hydrophobicity decoration (SHD [45]), the polymer scaling exponent, *ν*, derived from SHD and SCD [45], and the degree of aromatic clustering, which correlates with the material properties of protein assemblies [46]. Despite starting from disparate positions in clustering–*ν* space (Fig. S4), all replicas converged to similar values, showing that these structural features are co-selected with high fitness. Along the evolutionary trajectories, replicas 2 and 3 converged toward *ν* ∼ 0.5 – indicating selection for ideal chain behavior. Replica 1’s higher value (*ν* ∼ 0.6) suggests a marginally more expanded ensemble. This co-selection suggests that optimal lyotropic phase formation requires a balance of aromatic interactions (controlled by clustering) and appropriate chain flexibility (reflected in *ν*).

### 2.3 Structural and morphological features of peptide lyotropic phases

Having established convergence on a single molecular grammar for lyotropic phase formation – characterized by a sticker-spacer like architecture, aromatic clustering, specific amino acid preferences, and ideal chain behavior – we next ask what molecular level structural information this solution space encodes. To this end, we conduct a detailed structural and dynamical characterization of the lyotropic phase formed by a representative elite peptide from the optimal solution space. By the final iteration, the selected peptide [ENSFPFFN]_3_ had an average fitness score of −0.24 and ten repeats. These repeated evaluations both increase confidence in the fitness estimate and sustain the sequence’s genetic influence across subsequent iterations (see Methods). Hence, the peptide represents a well-validated solution within the optimized ensemble and serves as a prototypical example of the identified molecular grammar. We postulate that characterization of this peptide provides insight into the structural organization exhibited by the entire space of ‘elite sequences’. Indeed, consistent with this, the behavior observed for [ENSFPFFN]_3_ is closely mirrored by that of another independently selected candidate sequence [NQSGFFFS]_3_, shown in Fig. S5. In the following, we base our analysis on a 1 *µs* simulation of a 32×32 nm membrane consisting of 900 peptide molecules (Fig. 2A), selected to minimize finite-size boundary effects [47]. First, we analyze how the lamellar mesostructure emerges from collective single chain conformations. Secondly, we investigate the dynamics of chains within the membrane to determine whether this indeed reflects the liquid nature of a lyotropic phase.

**Figure 2.**
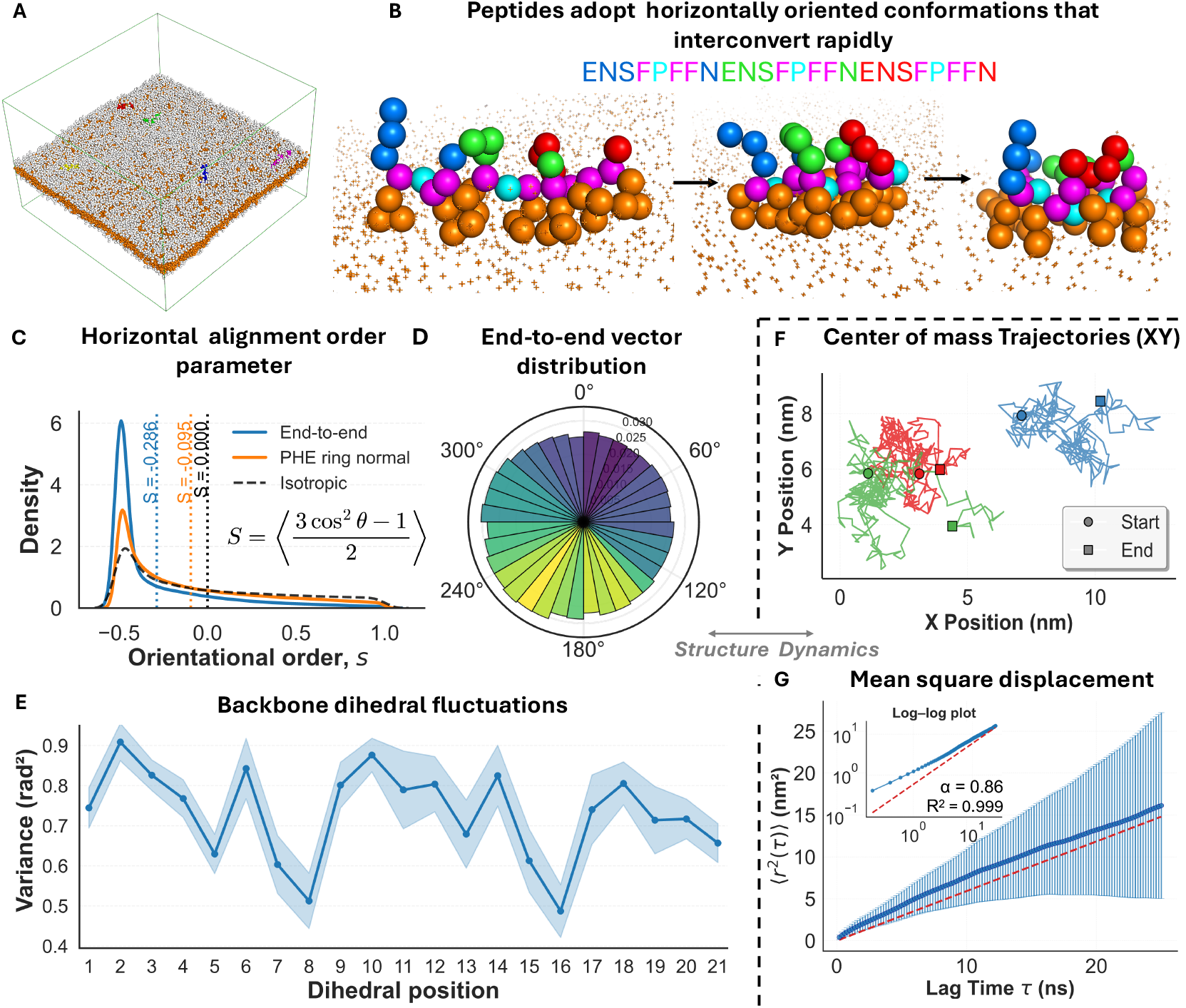
Structural and dynamic analysis of a coarse-grained lyotropic protein phase. (A) Bead representation of a simulated 32 × 32 nm membrane formed by 900 copies of [*ENSFPFFN*]_3_. Hydrophobic residues (orange) and hydrophilic residues (white). Five chains shown in different colors highlight horizontal embedding within the membrane. (B) Three conformations of the same membrane peptide at three different instances along the 1*µs* simulation trajectory. Backbone beads are colored according to the sequence displayed above; aromatic Phe sidechains are shown as orange beads. The chain adopts a horizontal conformation with phenylalanine side chains inserted into the hydrophobic core (orange crosses) while the backbone explores different conformations, from extended (left) to compact (right). (C) Distributions of the chain-wise orientational order parameter, *s*. The ensemble average ⟨*s*⟩ = *S* measures peptide alignment with respect to the membrane normal across all chains, time averaged to increase statistics. Vertical dotted lines indicate the *S* values for *θ*_E2E_, *θ*_Phe_, and an isotropic distribution, while solid lines show the corresponding *s* distributions. (D) Rose diagram of peptide chain end-to-end vectors projected onto the *x*–*y* plane. The angular distribution is approximately isotropic, indicating no preferred alignment. The radial extent of each bin corresponds to the probability that a chain within the ensemble is oriented along the angular range of that bin. The color of each sector encodes the mean end-to-end distance of chains in the bin, ranging from 17.5 Å (dark blue) to 19.5 Å (yellow). (E) Variance of dihedral angles along the peptide backbone reveals a periodic pattern where angles between repeat segments are most constrained (lowest variance). (F) Three representative peptide center-of-mass trajectories in the *x*–*y* plane. (G) Mean square displacement plot with log-log inset reveals normal fluid behavior with weak subdiffusion (*α* = 0.86) in two dimensions.

The repetitive multiblock sticker-spacer architecture drives peptides to adopt horizontal orientations within the membrane, lying flat with respect to the membrane normal (Fig. 2B). In this conformation, highly conserved phenylalanine residues organize across the bilayer with their benzene rings oriented towards the membrane center, forming a network of hydrophobic interactions which collectively form the hydrophobic core. The average geometric alignment of peptide chains with respect to the membrane normal was quantified via the order parameter *S* = ⟨*s*⟩, *i*.*e*. an ensemble average over all peptides and time. We calculated 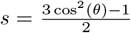 for two types of angles relative to the membrane normal: 1) *θ*_E2E_ – the angle between the peptide end-to-end vector and the membrane normal (z-axis), and 2) *θ*_Phe_ – the angle between the phenylalanine ring normal and the z-axis (Fig. 2C). Here, *S* = 0 corresponds to an isotropic angular distribution, positive values of *S* (0 *< S* ≤ 1) indicate preferential *parallel* alignment, and negative values ( − 0.5 ≤ *S <* 0) indicate preferential *perpendicular* alignment. The magnitude of *S* reflects the strength of the orientational order. Using this definition, we obtain 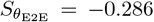 and 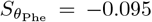. While 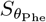 indicates no strong orientational preference for phenylalanine ring groups, the value of 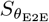 confirms that chains are systematically aligned horizontal to the membrane interface, which is also the only effective restriction to the peptide conformations. Otherwise, peptides adopt a broad array of conformations throughout the simulation trajectory. Single peptide tracking reveals exploration of a broad conformational ensemble, with an average end-to-end distance of 18.64 ± 8.18 Å (Fig. 2D). The normal distribution of end-to-end distances and the isotropic distribution of end-to-end vector directions in the membrane plane (Fig. 2D), coupled to the values for the orientational order parameter S, demonstrate that peptides are oriented with phenylalanine residues directed toward the membrane center, but not positionally registered, providing direct evidence for fluidity and lyotropic phase organization.

To probe the liquid nature of the membrane further, we conducted a mean squared displacement analysis by tracking the motion of all peptide centers of mass along the trajectory (Fig. 2F,G). A plot of lag time versus mean squared displacement reveals a linear relationship, with a diffusion coefficient of 0.1266 ± 0.0009 nm^2^/ns and a diffusion exponent *α* = 0.86 (fit *R*^2^ = 0.99) derived from the loglog plot (Fig. 2G inset plot), indicating weak subdiffusion in the membrane plane. This behaviour is consistent with known dynamics in the two-dimensional environment of lipid membranes, where transient interactions and local heterogeneity commonly give rise to subdiffusive motion [48, 49]. Having established liquid-like behavior of peptides within the membrane plane, we next extract peptide ensemble information related to the out-of-plane mechanical response, quantified in terms of its bending rigidity, *κ*. Analysis of the two-dimensional power spectrum of membrane height fluctuations yields *κ* = 7.9 ± 0.9 *k*_B_*T* for [ENSFPFFN]_3_ (Fig. S6), a value indicative of a flexible membrane that can readily accommodate curvature.

Molecular analysis also revealed position-specific conformational properties that correlate directly with functional requirements. Backbone pseudo-dihedral analysis (where the pseudo-dihedral is the angle between 4 consecutive backbone groups in the Martini force field) demonstrated clear periodicity matching the 8-amino acid repeat structure, with positions 5, 7, and 8 (and equivalent positions in each repeat segment) showing the lowest flexibility (Fig. 2E). These constrained dihedrals correspond to critical structural elements in the peptide: dihedral 5 (P**FF**N) spans the phenylalanine core region, while dihedrals 7 (F**NE**N) and 8 (N**EN**S) are those that define the inter-repeat hydrophilic spacer region. The reduced flexibility at these positions is an emergent property of membrane packing: as hydrophobic residues assemble to form the core, the developing lamellar structure constrains the dihedral angles to values consistent with the mesoscopic structure of the lyotropic phase. Consequently, limited dihedral fluctuations within the phenylalanine core (dihedral 5) maintain the positioning of hydrophobic residues that stabilize the membrane, while small fluctuations in dihedral 7 and 8 help to preserve the global peptide architecture. By contrast, the remaining backbone regions retain greater flexibility, meaning that they can undergo a greater degree of conformational adaptation whilst not disrupting lamellar organization.

Analysis of phenylalanine residue organization revealed remarkable stability through cooperative solvation effects. Despite high individual peptide mobility, 97.5% of all phenylalanine residues remained within the membrane core region throughout the simulation. Each phenylalanine residue maintained an average of 3.84 ± 1.07 hydrophobic contacts (*<*7 Å cutoff), providing sufficient solvation for core stability without excessive rigidity. Furthermore, across the entire membrane structure, a mean of 8.95 ± 2.02 of a total 8100 phenylalanine residues entered or left the core region per frame, showing that the membrane core composition remains remarkably stable. Individual peptides can thus undergo conformational changes and diffusive motion while the collective maintains the essential hydrophobic environment required for membrane function.

A prominent feature observed across the evolved sequences is the enrichment of proline and glycine residues at positions flanking at least one phenylalanine sticker. This observation is quite intriguing when considered within the context of the Martini coarse-grained force field. In this representation, both proline and glycine are distinguished from other amino acids solely by their small backbone beads, while their characteristic chemical properties – proline’s conformational rigidity due to its cyclic structure and glycine’s exceptional flexibility from the absence of a side chain – are not explicitly encoded through dihedral constraints or other chemical detail. Hence, we propose that the evolutionary selection for these residues reflects a geometric optimization: the reduced steric bulk of small backbone beads minimizes interference with the critical phenylalanine-phenylalanine interactions that stabilize the hydrophobic membrane core. This enables tighter intermolecular packing and stronger hydrophobic contacts essential for maintaining membrane integrity.

#### 2.3.1 Tuning solvent quality reveals the morphological space of lyotropic phases

Molecules forming lyotropic phases in dilute aqueous solution adopt morphologies determined primarily by molecular packing (Fig. 3A). In this framework, changes in solvent quality alter the relative effective volumes of hydrophilic and hydrophobic segments, shifting preferred packing and thus mesoscale structure. The resulting morphology is commonly rationalized by the packing parameter, which captures the balance between interfacial area, hydrophobic volume, and chain length [51].

Here, we analyze the phase behaviour of lyotropic phase forming peptides by tuning the solvent quality for a set phase (lamellar). We emphasize that the lyotropic phases observed are not artificially stabilized by periodic boundary conditions, as the simulation box was chosen to be sufficiently larger than the dimensions of the assembled structure. In Martini 3, the parameter *λ* is used to modulate solvent quality and uniformly scales protein-water interactions in order to reproduce experimental radii of gyration for IDPs [52]. As solvent quality improves (i.e., higher *λ*), the hydrophilic segments swell relative to the hydrophobic ones, increasing the effective hydrophilic–hydrophobic volume ratio. As predicted by the aforementioned theory, when the solvent quality is systematically varied in our isotropic pressure simulations, the relative hydrophobic and hydrophilic volume is modulated, giving rise to a different optimal packing and therefore a change in morphology. The morphologies accessed upon changing *λ* reproduce the expected morphological sequence for lyotropic phases. At the standard value of *λ* = 1, [ENSFPFFN]_3_ forms a bicelle, which in lyotropic phase theory corresponds to *C*_*pp*_ ≈ 1, or a cylindrical packing geometry. This is consistent with the lamellar phase identified during EvoMD optimization. At a slightly increased *λ* = 1.04, a phase boundary is encountered. Here the chemical potential of peptides in both bicelle and worm-like micelle morphologies are roughly equal, leading to a dynamic equilibrium between the two morphologies. At *λ* = 1.05, the worm-like micelle is stable, and at *λ* = 1.06, the system transitions to form spherical micelles. As the value of *λ* is increased from this point, the average number of peptides per micelle, or micelle aggregation number, decreases, until at *λ* = 1.09, the solution is in a disordered state, with all peptides in their fully solvated, monomeric form. This systematic progression through bicelle, worm-like micelle, cylindrical micelle, and disordered states upon varying solvent quality confirms that these peptide phases are governed by the classical packing parameter concept and genuinely lyotropic in nature.

**Figure 3.**
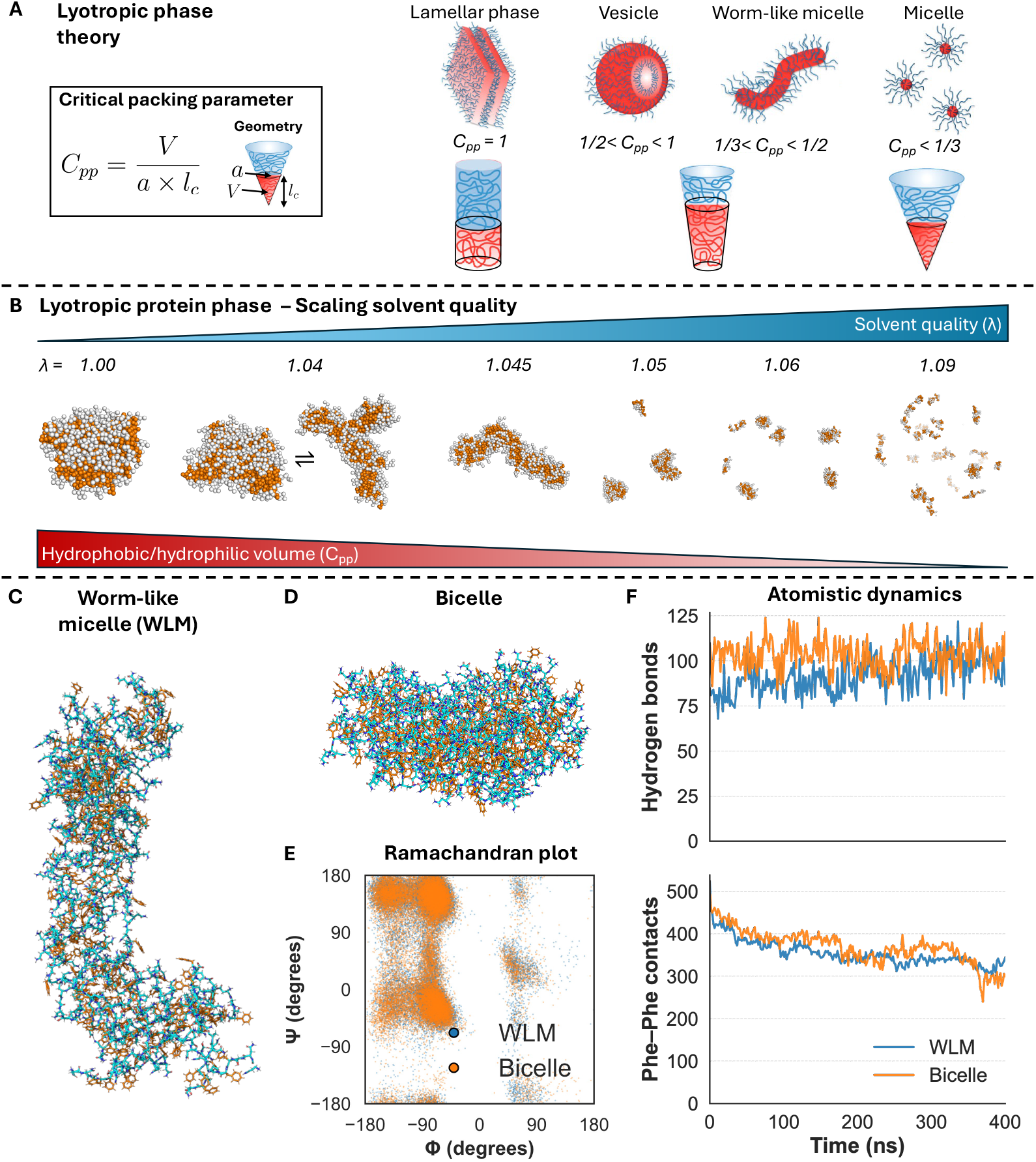
Analysis of the lyotropic phase morphology. (A) Schematic showing the general effect of the critical packing parameter (*C*_*pp*_) on lyotropic phase morphology. *C*_*pp*_ = *V/*(*a* × *l*_*c*_), where *a* is the interfacial cross-sectional area between hydrophilic and hydrophobic segments, *V* is the hydrophobic segment volume, and *l*_*c*_ is the hydrophobic segment length. These parameters set the preferred interfacial curvature, giving rise to morphological transitions from lamellae to vesicles to worm-like micelles to micelles with decreasing *C*_*pp*_ *(adapted from Nunes et al. [50]*). (B) A simulated example evolves from lamellar to worm-like micelles to cylindrical micelles to disordered as solvent quality increases (increasing *λ*, scaling the protein-water interactions). This corresponds to a decrease in *C*_*pp*_ as outlined in panel A. (C) Atomistic simulations performed on (i) WLM and (ii) bicelle phases, testing their stability in all-atom MD. Initial structures were generated by backmapping equilibrated Martini coarse-grained configurations from panel B. Phenylalanine sidechains (orange) and backbone/sidechain carbons (cyan) are shown as a licorice representation. (D) Ramachandran plot of *ϕ* and *ψ* angles across 400 ns for all chains in both systems. No preference for *β*-sheet versus *α*-helix, consistent with an amorphous phase. (E) Hydrogen bonds (top) and Phe-Phe contacts (bottom) versus simulation time for both systems.

#### 2.3.2 From Coarse-Grained to Atomistic: Explicit Directional Interactions

To gain insight into the role of directional interactions in the late stages of lyotropic peptide assembly, we performed a series of atomistic molecular dynamics simulations. Lyotropic phases at the end of Martini coarse-grained simulations were backmapped to the atomistic level and simulated for 400 ns using the a99SB-disp force field, which is optimized to reproduce conformational ensembles of both disordered and globular proteins [53]. Specifically, we sought to determine whether the lyotropic structures are stable when including explicit directional interactions, and whether a structurally feasible set of peptide conformations can yield such a phase. As the underlying free-energy landscape depends on the resolution of the molecular representation, the thermodynamic state may not be exactly preserved and phase boundaries could shift, meaning that a given sequence may favor an adjacent equilibrium morphology in atomistic resolution compared to CG resolution. To accommodate such an effect, we backmapped two CG morphologies for the sequence [NNSFFPFN]_3_: the worm-like micelle (WLM) and the bicelle (Fig. 3C,D), and chose a neutral system to avoid artifacts arising from the differential treatment of electrostatics in the CG and atomistic simulations. We find that both backmapped structures converge to stable interaction profiles, as quantified by hydrogen-bond and Phe–Phe contact counts, over 400 ns of simulation (Fig. 3F). Analyses of backbone dihedral angles over the course of each simulation additionally reveal a heterogenous distribution of secondary structures (*ϕ*/*ψ* pairs) for both morphologies, consisting of *β*-sheet structural elements as well as right- and left-handed *α*-helices (Fig. 3E). This shows that the lyotropic structures are composed of peptide conformations that are structurally permitted at the atomistic level. Overall, these results confirm the feasibility of the lyotropic phase in the presence of explicit directional interactions.

### 2.4 A shared molecular grammar links lyotropic phases and amyloid fibrils

Sequences that form biomolecular condensates are increasingly recognized to also access amyloid states, either from the dilute phase or through condensate maturation, reflecting an overlap in their underlying sequence grammars [30]. Optimization for lyotropic-phase formation yields sequences sharing key features with prion-like domains and LARKS motifs, which are characteristic of both condensate-forming proteins and amyloid fibrils. To further elaborate on their relation, we employed structure prediction with AlphaFold 3 [54] and assessed whether peptides exhibiting lyotropic molecular grammar are predicted to adopt amyloid-like architectures as structural end points. Although application of AlphaFold 3 for structure prediction in IDPs is in its infancy and raises conceptual issues [55], we expect that it provides useful insight for semi-crystalline aggregated states [54]. Indeed, AlphaFold 3 accurately predicts the molecular geometries of decameric (10-mer) assemblies of experimentally characterized LARKS sequences – SYSGYS (PDB ID: 6BWZ), GFGNFGTS (PDB ID: 6BZM), GYNGFG (PDB ID: 6BXX), SYSSYGQS (PDB ID: 6BXV), and STGGYG (PDB ID: 6BZP) – with an average monomer C*α* RMSD of 1.03 Å and a whole-assembly C*α* RMSD of 2.25 Å relative to PDB structures (Fig. S9). AlphaFold 3 achieved similarly high accuracy for the canonical amyloid steric zipper A*β*_16–21_, yielding monomer and assembly C*α* RMSDs of 0.36 Å and 0.441 Å, respectively (Fig. 4, Fig. S9, Table S1). These results indicate that AlphaFold 3 does provide reasonable predictions of amyloid-like structural end points for sequences within this molecular grammar. We then applied AlphaFold 3 to decameric assemblies of sequences drawn from the lyotropic-phase forming sequence space – ENSFPFFN, NQNGFPFD, ENGFPFFN, and IWWPANNG (from the `no PHE` run) – to assess the diversity of predicted amyloid-like architectures encoded within the lyotropic sequence landscape. Despite sequence variation, all predicted structures adopt amyloid-like architectures (Fig. 4), indicative of the existence of a shared molecular grammar among prion-like domains, LARKS, and our designed lyotropic phase-forming sequences. It provides additional evidence that amyloid-like architectures are a sequence encoded structural end point for lyotropic phase forming peptides.

**Figure 4.**
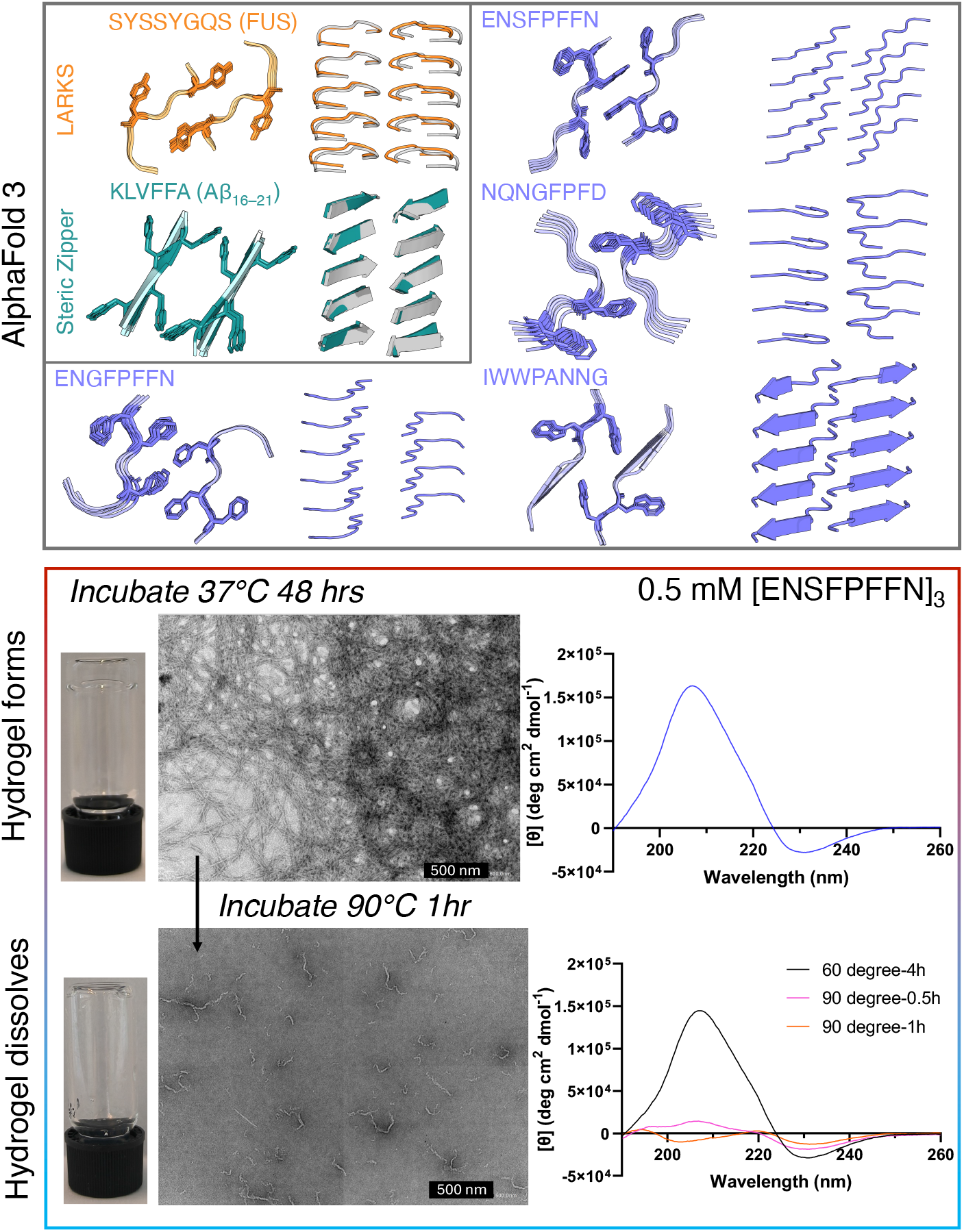
AlphaFold 3 predictions and experimental characterization of fibrillar end points. (Top) AlphaFold 3 predicted fibrillar architectures for selected lyotropic phase forming motifs, shown in purple. As an offset, predictions for previously determined, natural motifs; a canonical LARKS motif, SYSSYFQS (orange), and the steric zipper KLVFFA (blue). Overlays with X-ray crystal structures of SYSSYGQS and KLVFFA from the Protein Data Bank show close agreement between predicted and experimentally determined amyloid structures. (Bottom) Experimental: peptide [ENSFPFFN]_3_ (0.5 mM) was incubated in 0.1 mM NH_4_Cl (pH 7.6) at 37 ^*◦*^C for 48 h. Transmission electron microscopy (TEM) revealed a dense network of interconnected fibrils (scale bars, 500 nm). Hydrogel formation was confirmed by the inverted vial test. Circular dichroism spectroscopy indicated a *β*-sheet–rich secondary structure. Subsequent heating to 90 ^*◦*^C for 1 h disrupted the hydrogel and markedly reduced *β*-sheet content, consistent with fibril dissociation, while residual fibrillar structures remained detectable by TEM.

Complementing these ML predictions, we also examined the assembly behavior of [ENSFPFFN]_3_ under *in vitro* conditions. Circular dichroism (CD) and transmission electron microscopy (TEM) were used to characterize the structural evolution, and macroscale gelation was assessed with inverted-vial assays. Under the *in vitro* conditions, the experimental observables primarily report on equilibrium end-point morphologies rather than structures that we predict to arise during the initial stages of self-assembly (see Discussion). The transient early stage development of lyotropic structure is apparently elusive to the considered techniques, meaning that the results report exclusively on aging into the equilibrium end-point morphologies. CD spectroscopy shows that 0.5 mM [ENSFPFFN]_3_ rapidly adopts *β*-rich structure at 37 ^*◦*^C in buffered aqueous solution (pH 7.6, 0.1 mM NH_4_Cl). Even at the initial 0 h timepoint (measured several minutes after sample preparation), the CD spectrum exhibits a weak but discernible *β*-sheet signature, characterized by a broad maximum near 210 nm and a depression near 230 nm (Fig. S11). The amplitude of these features increases substantially by 4 h, consistent with the growth of *β*-structured assemblies. The presence of detectable secondary structure at 0 h suggests that amyloid-competent oligomers form within minutes of solubilization, with progressive formation of *β*-rich assemblies over the first few hours. By 48 h, the spectral shape remains similar but increases further in intensity (Fig. 4), indicating continued accumulation or consolidation of *β*-rich fibrillar material. An inverted-vial assay confirms that by 48 h the peptide forms a self-supporting hydrogel. Transmission electron microscopy (TEM) imaging of the gel reveals a dense fibrillar network with morphology characteristic of amyloid-based hydrogels, including long, unbranched filaments and cross-linked bundles. Thermal stability assays further demonstrate that these assemblies possess the characteristic robustness of amyloid structure. Heating the gel to 60 ^*◦*^C for 4 h leaves the CD signature largely intact (Fig. S11), indicating preservation of the underlying *β*-sheet architecture. In contrast, exposure to 90 ^*◦*^C induces melting: the gel liquefies, the *β*-signal decreases markedly within 30 min, and after 1 h the spectrum shifts to display two minima (near 205 nm and ∼ 210 nm), consistent with substantial secondary-structure reorganization (Fig. 4). Collectively, these data demonstrate that [ENSFPFFN]_3_ readily forms amyloid fibrils under the conditions used in our experimental setup, in line with the expected aging behavior. More broadly, these findings suggest that the self-assembled structure of a peptide, whether a lyotropic phase or an amyloid state, emerges from kinetic control of the assembly pathway. Tuning the properties of amphiphilic sticker motifs reshapes this kinetic landscape, as discussed in the following section.

## 3 Discussion

Over the past years, it has become increasingly clear that the gap between one-dimensional, paracrystalline amyloid fibrils and three-dimensional liquid droplets is narrower than once thought [30]. Biomolecular condensates are now recognized as metastable, out-of-equilibrium systems with material properties that are regulated through various mechanisms [56]. In the absence of regulation, condensates progress toward equilibrium through molecular events that favor the global thermodynamic minimum – amyloid fibrils. Alternatively, the dense phase can mature into a kinetically trapped gel state [56]. The balance among these outcomes, sustained by various cellular processes, is tightly linked to condensate functionality.

Despite progress, our understanding of the molecular mechanisms that govern these mesoscopic phase transitions remains in its infancy. We hypothesize that canonical sticker–sticker interactions between prion-like motifs in pron-like domain (PLD)-containing condensates give rise, over time, to disordered clusters that constitute lyotropic phases and provide nucleation-competent microenvironments for amyloid fibril formation. To test this hypothesis, we employed a *de novo* inverse design approach, coupling genetic algorithms with molecular dynamics simulations to determine the space of protein sequences capable of forming lyotropic phases.

The significant finding is that in searching for lyotropic phases, this design approach – which does not presuppose any molecular grammar – independently converges on the molecular grammar of PLDs that are known to form an array of both condensates and amyloids [39]. Sequence logos across all three replicas reveal strong enrichment for phenylalanine, tryptophan, proline, glycine, asparagine, serine, glutamine, and alanine (Fig. 1E), all (with the exception of proline) known to occur prevelantly in PLDs [57]. This enrichment pattern is not arbitrary – the higher-order architecture of our sequences matches closely to that of LARKS (Low-complexity Aromatic-Rich Kinked Segments) [21, 23], exhibiting aromatic clusters flanked by glycine and proline residues, where G and P permit the formation of kinks that prevent rigid beta-sheet formation while allowing for reversible self-assembly. A crucial finding is that a short lyotropic motif of only 8 amino acids, on the same length scale as LARKS like stickers which are 6 to 8 amino acids in length, is sufficient to drive the formation of lyotropic assemblies. The molecular grammar of the 8 amino acid repeat length encodes chain conformations where backbones are oriented perpendicular to the membrane normal, with aromatic stickers pointing towards the center to form the hydrophobic core. This results in a chain conformation that is characterized by the fact that upon microphase separation [58–60], both peptide termini end up being situated on the same side of the membrane. This implies that in the conditions where lyotropic phase formation is favored, a lyotropic sticker motif embedded between two spacer sequences (much like LARKS, which are interspersed throughout IDPs such as FUS) can form a homotypic interaction (Fig. 5) with an equivalent sticker motif on another chain whilst maintaining the topological requirements for phase formation. This is only possible in multiblock chain topologies and not in a classical di-block copolymer, where the amphiphillic group would need to be present at the chain terminus to allow for a lyotropic phase to form.

**Figure 5.**
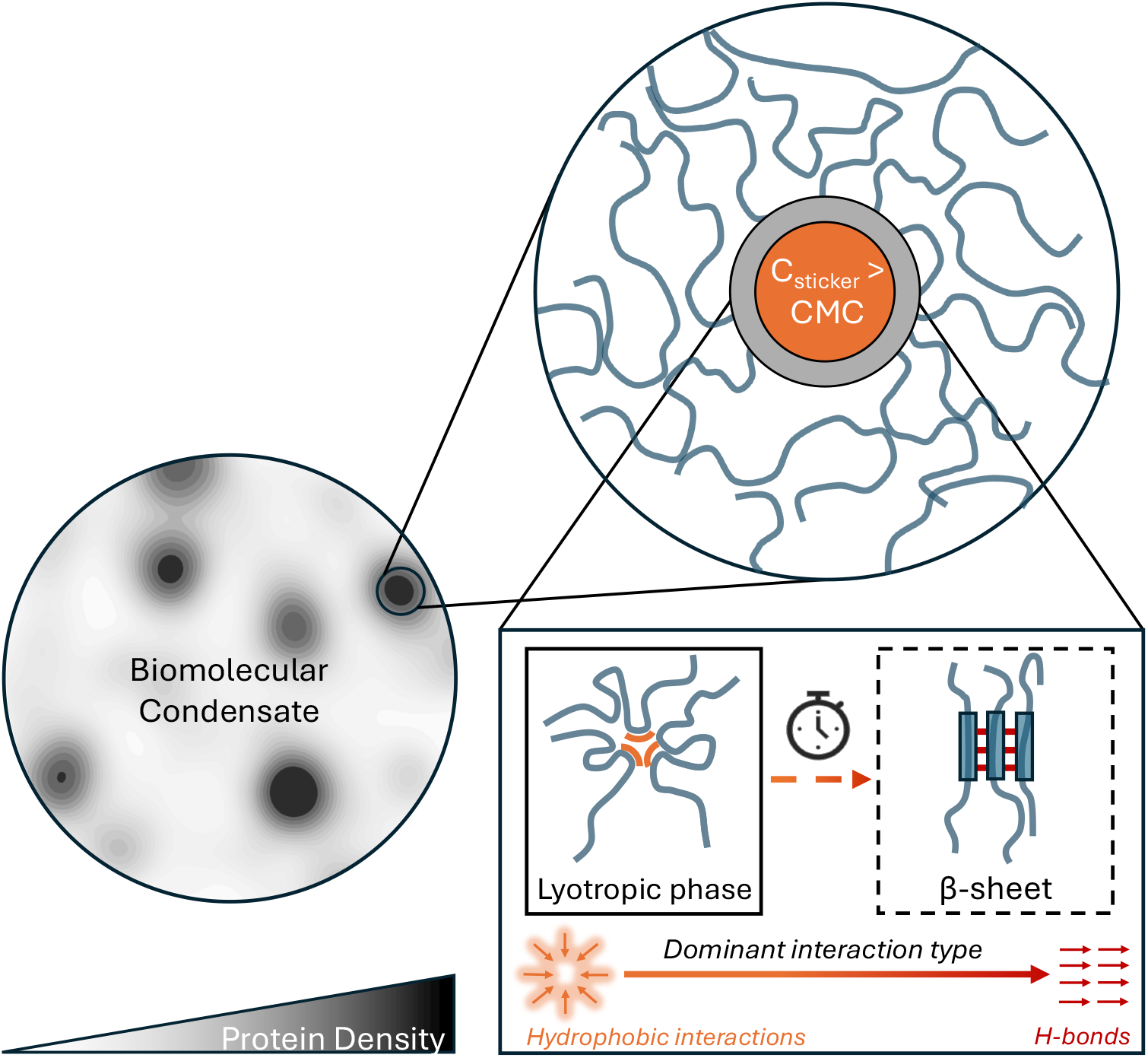
Schematic depiction of lyotropic phase formation in aging biomolecular condensates. Protein density fluctuates within the condensate, and in high-density regions the local sticker concentration exceeds the critical micelle concentration (CMC). This drives the formation of lyotropic phases. Within lyotropic phases, stickers engage in cooperative hydrophobic interactions, which do not require precise alignment. This low geometric specificity is represented by soft, glowing orange interaction arrows, indicating that stickers can shift relative positions while maintaining favorable contacts. Over time, chain segments within the lyotropic phase may rearrange such that they are increasingly stabilized by a semi-crystalline network of hydrogen bonding interactions. This causes local *β* sheet structures to form. The directionality of these interactions is represented by the red arrows. In most cases, due to the high viscosity and poor solvation of the condensate microenvironment, these structures remain kinetically trapped and are unable to reorganize into extended amyloid fibrils.

The strong overlap between the lyotropic and PLD molecular grammar signifies the biological relevance of our computationally designed peptides. The grammar directly connects to the canonical amyloidogenic motif of amyloid-*β*_16−21_ – KLVFFA [61], with the FF motif being the most strongly occuring 2-mer in the lyotropic sequence space (Fig. S3)[62]. Indeed, there is a remarkable resemblance between our micellar assemblies (Fig. 3B) and the stable, globular and disordered oligomers yielded by the 2-step nucleation mechanism of A*β*_16−21_ oligomerization [63–65]. This mechanism occurs when the formation of hydrophobic interactions between Phe residues is favoured over the formation of hydrogen bonds. It reflects the nature of our simulations, where the Martini force field employed only represents isotropic interactions and lacks explicit directional interactions such as hydrogen bonding [66]. Therefore by design, our simulations provide insight into the (early) stages of protein self-assembly processes where hydrophobic interactions energetically dominate over directional hydrogen bonds [67– 70]. Additionally, analysis of aromatic clustering and hydrophobic content places our elite sequences in precisely the same region as Velo1-PLD [46](Fig. S12), a prion-like domain known to form reversible, solid biomolecular condensates enriched in amyloid-like fibers that scaffold Balbiani bodies [71, 72]. It suggests that this region of sequence space possesses the features required for the formation of a reversible lyotropic phase, perhaps holding promise as a model for understanding the elusive phase separation of the Balbani body.

Experimentally, the structural characteristics of our designed peptide [ENSFPFFN]_3_ directly reflect the properties expected for a peptide with the same aromatic packing and hydrophobic content as Velo1PLD [46, 73], and the FF motif of A*β*_16−21_[62]. We find our system readily forms amyloid fibrils that gelate into a network structure, undergoing melting at 90°C. Though this is a higher melting point than measured for a LARKS gel, we hypothesise that the length and imposed symmetry of our chain lends itself to self-templating and therefore the formation of more stable amyloid-like fibers [74]. Also, experimental conditions such as the peptide concentration and nature of the buffer affect the formation kinetics. To shed light on this, beyond the range of this study, further investigations could consider individual 8AA lyotropic motifs, or more heterotypic systems consisting of different identified lyotropic motifs embedded between long chains of stickers. Additionally, modifying solvent conditions to those that disfavor directional hydrogen bonding or enhance the hydrophobic effect would enable the investigation of 2SN like disordered intermediates prior to amyloid conversion [75–78].

Beyond the conserved aromatic stickers, our evolutionary optimization reveals that surrounding spacer residues exert fine control over lyotropic morphology. While the hydrophobic sticker block (particularly FF) remains highly conserved, single point mutations in spacer regions can shift the system between different lyotropic morphologies (Fig. S7,S8). For example, polar-to-charged transitions (N to E) increase the relative volume of the hydrophilic block, causing lamellar-to-worm-like micelle transitions. This fine-tuning capacity suggests that natural PLDs could modulate condensate material states through spacer variations while preserving core sticker function [79].

### 3.1 Lyotropic phases as key intermediates in condensate maturation

Focusing on kinetic control and aging, we propose that the lyotropic phase represents a key nanoscopic structural element that promotes viscoelastic maturation in condensates. Lyotropic assemblies act as physical cross-links [60, 80–82] that occupy an intermediate energetic and kinetic regime between highly transient sticker–sticker contacts and stable, crystalline amyloid phases. The interior of condensates, we argue, provides an environment that strongly favors their formation.

Lyotropic assemblies form via a nucleation process that is governed by the critical micelle concentration (CMC) [83]. Below the CMC, aromatic and hydrophobic motifs simply act as transient dimeric stickers; above it, they cooperatively condense into lyotropic microphases. Because lysolipids of comparable size exhibit micromolar CMCs [84], the markedly higher protein concentrations inside condensates – reaching the millimolar range ( ∼ 2 mM for FUS) [85, 86] – place these systems firmly in the lyotropic regime. This being said, although exceeding the CMC is a necessary condition for lyotropic phases to form, it is not sufficient on its own. The CMC is defined under equilibrium conditions, whereas biomolecular condensates are intrinsically non-equilibrium systems. As a result, whether a lyotropic equilibrium can be attained within condensates depends on additional features of the internal condensate environment that govern kinetic accessibility, particularly protein hydration and conformational dynamics.

At millimolar concentrations, proteins experience high local crowding and are poorly solvated, with overlapping hydration shells. Terahertz spectroscopy shows that, as a result, weakly bound water molecules associated with hydrophobic residues become destabilized and expelled in phase-separated FUS droplets [85]; an effect that in turn increases the overall entropy of the system and provides a significant entropic driving force for LLPS. Consequently, nondirectional hydrophobic interactions are strongly favored within the condensate interior, increasing the probability of lyotropic phase formation. In parallel, and from the perspective of protein conformational dynamics, chains in the dense condensate interior cannot readily reorient or slide because they have a high internal friction [87]. This is because chains are constrained by multiple weak sticker–sticker interactions and by topological factors, such as the network dimensions and entanglements [88] that are distributed along their contour. Large-scale rearrangements therefore require coordinated disengagement of many local interactions and are significantly slowed down [89]. We propose that lyotropic assemblies, which form through nondirectional hydrophobic collapse, are thus much more likely to *initially* appear than ordered *β*-structures, whose formation requires slow and highly coordinated chain rearrangements. Subsequent progression from lyotropic assemblies toward *β*-rich or crystalline states then depends on whether spacer solvation permits such rearrangements [90]. Notably, *β*-sheet and amyloid structures tend to appear preferentially at interfaces during aging, where heterogeneous nucleation relaxes many of the constraints that frustrate ordering in the core. At interfaces, partial dehydration, reduced dimensionality, and altered chain orientation lower the entropic cost of alignment and therefore increase the probability of amyloid competent chain arrangements [25, 27, 28, 76, 91].

Because lyotropic assemblies are stabilized primarily through cooperative hydrophobic collapse, perturbations to aromatic packing geometry should directly modulate their stability. Consistent with this expectation, our results indicate that tyrosine may reduce susceptibility to pathological solidification. Substituting phenylalanine with tyrosine in the lyotropic phase–forming peptide [ENSFPFFN]_3_ to generate [ENSYPFYN]_3_ demonstrates that, although a lyotropic phase still forms, the phenolic hydroxyl disrupts efficient packing within the hydrophobic core, producing a jagged and geometrically frustrated interface (Fig. S13). Thus, even when the CMC is exceeded and condensation occurs, the free-energy minimum of the tyrosine-rich lyotropic assembly is likely shallower than that of its phenylalanine-rich counterpart. By penalizing tight hydrophobic packing while introducing, through its hydroxyl group, additional polar and hydrogen-bonding interactions that compete with aromatic packing, tyrosine shifts the equilibrium away from dense lyotropic assemblies. Although alternative pathways to amyloid formation may exist, densely packed lyotropic phases provide concentrated, structurally biased environments that seed *β*-sheet and amyloid fibril formation (Fig. 5). Destabilizing this assembly reduces the likelihood of accessing this pathway, suggesting that tyrosine enrichment in naturally occurring PLDs may reflect a dual functional advantage – its well-established role as a strong sticker promoting dynamic multivalent interactions [92, 93] and its apparent capacity, demonstrated here, to limit pathological solidification. This interpretation is further supported by bioinformatic analyses indicating preferential enrichment of phenylalanine in more solid-like condensates and of tyrosine in more liquid-like assemblies [94].

Lyotropic phases also provide a physical interpretation for the nanoscale architecture predicted by phase separation coupled to percolation (PSCP) [95]. In the PSCP model, associative polymers above both the saturation and percolation thresholds (*C*_*sat*_ *< C*_*perc*_ *< C*_*dense*_) form percolated networks with small-world topology [28]. These networks comprise dense molecular clusters (“cliques”) connected by extended chains (“hubs”) that engage in transient interactions to ensure global connectivity [28, 96]. Coarse-grained molecular dynamics simulations show that cliques contain compact, quasi-spherical chains with confined yet rapid local motion, forming structures with long lifetimes, whereas hubs contain more extended, dynamic chains that bridge distant cliques [96]. Experiments support this architecture: nanoscale cliques with distinct environments have been observed in FUS [97] and A1-LCD condensates [98], and significant spatial heterogeneity – with dense “liquid-like” and dilute “gas-like” regions – has been documented in multicomponent systems [99]. We propose that these cliques correspond to regions that provide the microenvironment necessary to stabilize lyotropic assemblies – in particular, regions where high chain concentration, hydration shell overlap, and restricted mobility jointly favor nondirectional hydrophobic association—whereas hubs reflect environments where chains remain sufficiently extended and mobile to percolate the network.

The resulting physical form of the network of cliques and hubs generates multiscale spatiotemporal and thermokinetic inhomogeneities that dictate condensate rheology. Over time, these inhomogeneities evolve, causing condensates to transition from Maxwell fluids to materials dominated by elastic responses (G^*′*^ *>* G^*′′*^), exhibiting glassy aging reminiscent of microgels [24, 90]. This transition is not typically accompanied by the appearance of crystalline *β*-rich structures: in experiment, aging condensates often remain amorphous, lack ThT staining, and show no evidence of fibrillar ordering [24, 86, 90]. We note that this behavior parallels polymer gel systems in which structural inhomogeneities arising from critical fluctuations and microphase separation characterize the onset of gelation [100].

Within this context, lyotropic phases emerge as key intermediates that help drive the viscoelastic maturation of condensates. Most lyotropic assemblies remain amorphous and reversible. However, their intermediate stability and inherent high local protein concentration render them as hotspots at which cooperative alignment events may occur, enabling undesired localized progression toward *β*-rich or amyloid-like states under specific conditions. Thus, lyotropic phases unify the thermodynamics of sticker association with the emergent network architecture and viscoelastic maturation of condensates. As an exciting prospect, they may represent tractable targets for interventions aimed at preventing pathological aging before irreversible *β*-structured or amorphous vitrified states emerge.

Together, our findings suggest that lyotropic microphase formation is an intrinsic feature of prion-like sticker grammars, whose inherent amyloid-seeding potential exerts evolutionary pressure on PLD sequence composition. These prion-like motifs evolved to form and stabilize condensates via transient, dimeric, hydrophobic, sticker-to-sticker interactions, all the while conserving protein solubility. This process requires motifs with balanced amphiphilicity to direct and shield hydrophobic interactions without collapsing completely. However, the nature of this solution space favors larger, thermodynamically stable, dynamic assemblies – lyotropic phases – that can seed amyloid fibrils as an unintended, pathological side effect over prolonged aging or stress.

## 4 Methods

### 4.1 Evolutionary Molecular Dynamics (Evo-MD) design framework

Peptide sequences were designed *de novo* using Evo-MD, an in-house physics-based inverse-design framework which we adaped from previous works [37, 101] to design lyotropic phase forming peptides. For Evo-MD–based optimization of lyotropic phases, we ran 20 iterations of the genetic algorithm, where 512 peptides were evaluated per iteration. In the initial iteration, each of the 512 sequences were constructed by sequentially selecting 8 amino acids at random to form an 8AA segement, and repeating this segment three times to produce a 24-residue peptide. To sample the relevant space of intrinsically disordered peptides, amino acids were drawn according to their frequencies in the DisProt database [102], and sequences were retained only if all residues were predicted to adopt coil secondary structure by the Porter5 secondary structure prediction tool [103].

For each peptide within the optimization procedure, we ran a coarse-grained self-assembly simulation of 25 peptides, and then analyzed the resulting simulation to quantify the ability of the peptide to form a lyotropic phase (the lyotropic fitness function is described below). Upon fitness evaluation, the top 64 scoring sequences were selected as parents for constructing the population of the next iteration. The top 32 sequences were classified as elites and passed to the next iteration without alteration, thereby contributing further to the genetic information available for recombination. Elite sequences were reevaluated at each iteration, with their reported fitness averaged over the current and all previous evaluations, yielding progressively more reliable fitness estimates. Sequences for each subsequent iteration were generated by uniform crossover between parents, followed by point mutations applied with probability 1*/N* per residue (*N* = 24). This selection–recombination–mutation cycle iteratively drove the population toward peptides that form lyotropic phases.

### 4.2 Evo-MD coarse-grained simulations and fitness evaluation

Coarse-grained MD simulations were performed with the Martini 3 force field [40] using GROMACS 2023.3 [104]. For each candidate peptide, a coarse-grained monomer was generated using martinize2 [105] and then assembled into a slab configuration containing 25 copies of the peptide. Systems were energy-minimized, equilibrated, and simulated for 400 ns in the semi-isotropic NPT ensemble at 300 K with Langevin dynamics. Temperature was maintained using the Langevin thermostat, and pressure was controlled at 1 bar using the standard semi-isotropic Berendsen coupling for Evo-MD [37, 101]. Nonbonded interactions followed standard Martini 3 settings, including a 1.1 nm cutoff for van der Waals and reaction-field electrostatics and the Verlet cutoff scheme for neighbor searching. Full simulation details are provided in the Supplementary Information.

To search for lamellar phases, we designed a fitness function that forces the emergence of orientational anisotropy along one axis of the dense phase. In our slab geometry, the long axis (which we here select as the z-axis, *L*_*Z*_ *>> L*_*X*_, *L*_*Y*_) serves as the direction along which ordered structures develop preferential molecular alignment, contrasting with the random orientations characteristic of isotropic liquid phases. The fitness function minimizes the spatial overlap between hydrophobic and hydrophilic amino acid residues within the system

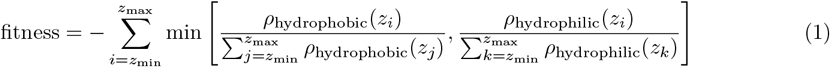

Here, *ρ*_hydrophobic_(*z*) and *ρ*_hydrophilic_(*z*), represent the number densities of hydrophobic and hydrophilic amino acids, respectively, at a given point *z* along the z-axis (See SI Methods for AA classification). The value of the fitness function ranges between −1 and 0, where −1 corresponds to the minimum fitness (disordered) and 0 corresponds to the maximum fitness (lamellar). All densities were computed along *z* using the GROMACS `gmx density` tool.

### 4.3 Large-scale membrane simulations

Large-scale membrane systems (32 × 32 nm, 900 peptide chains) were constructed with periodic copies of equilibrated slab configurations generated during the genetic algorithm optimization. These systems were simulated for 1 *µ*s in the semi-isotropic NPT ensemble at 300 K using the same thermostat and nonbonded parameters as in the Evo-MD runs, with pressure controlled using the Parrinello–Rahman barostat.

#### 4.3.1 Trajectory analysis

Trajectory analysis was performed using MDAnalysis [106]. We quantified peptide orientation, in-plane and out-of-plane nematic ordering, center-of-mass diffusion, backbone dihedral flexibility, hydrogen bonding, and aromatic–aromatic interactions using established geometric definitions. Aromatic contacts between phenylalanine residues were defined using a 7 Å centroid-to-centroid cutoff, consistent with established ranges for favorable Phe–Phe interactions [107]. Membrane bending rigidities were estimated from thermal undulations using a custom protocol which combines MDVoxelSegmentation [108] with an analysis framework adapted from Fowler *et al*. [109]. Complete analysis definitions, atom selections, and statistical procedures are described in the Supplementary Information.

### 4.4 Solvent-quality (*λ*-scaling) simulations

Solvent-quality modulation was performed using the *λ*-scaling protocol of Thomasen *et al*. [52], in which all peptide–water Lennard-Jones interactions are scaled by a factor *λ*. For each selected peptide, an equilibrated 25 peptide slab from the Evo-MD protocol was made whole, placed into a much larger 25 × 25 × 25 nm water box to remove boundary effects, and simulated for 500 ns in the isotropic NPT ensemble at 300 K. Temperature was controlled with the velocity-rescaling (V-rescale) thermostat, and pressure was maintained at 1 bar using isotropic Berendsen coupling. To enable direct comparison, all other parameters matched those used for the coarse-grained slab simulations. Full details of construction and equilibration are provided in the Supplementary Information.

### 4.5 All-atom validation

Representative coarse-grained assemblies were backmapped to atomistic resolution using ‘backward’ [110] and simulated using the a99SB-disp force field [53]. Backmapped systems were energy minimized, equilibrated sequentially in NVT and NPT ensembles, and simulated for 400 ns at 300 K in the NPT ensemble. Temperature was maintained using the V-rescale thermostat, and pressure was controlled using the Parrinello–Rahman barostat at 1 bar. Electrostatics were treated with the Particle Mesh Ewald method using a 1.0-nm real-space cutoff. Complete preparation, equilibration, restraint, and integration parameters are provided in the Supplementary Information.

### 4.6 AlphaFold 3 structure prediction

Peptide structures were predicted using AlphaFold 3 [54] with default parameters. Predicted models were compared to available experimentally determined structures deposited in the Protein Data Bank (PDB). Structural alignments were performed in PyMOL, and root-mean-square deviations (RMSDs) were calculated for both monomeric peptides and whole assemblies where applicable. RMSDs were computed separately for all atoms and for C*α* atoms. Model confidence was assessed using the mean pLDDT score, as well as the interfacial predicted TM-score (iPTM) and predicted TM-score (PTM) reported by AlphaFold.

### 4.7 Experimental materials and methods

#### 4.7.1 Materials

Fmoc-protected amino acids, ethyl cyano(hydroxyimino)acetate (Oxyma), nitrosobenzene, *N–N*^*′*^-diisopropylcarbodiimide (DIC), triisopropylsilane (TIS), and Formvar/Carbon-supported copper (200 mesh) TEM grids were purchased from Sigma-Aldrich. Trifluoroacetic acid (TFA), piperidine, dimethylformamide (DMF), and acetonitrile (MeCN) were purchased from Biosolve (Valkenswaard, The Netherlands). Diethyl ether (Et_2_O), dichloromethane (DCM), and methanol (MeOH) were supplied by Honeywell (Meppel, The Netherlands). Ammonium hydroxide (NH_4_OH) was purchased from ThermoFisher scientific. Milli-Q water was obtained using a PURELAB flex water purification systems (ELGA LabWater, UK). TentaGel^®^ S RAM resin was purchased from Rapp Polymer GmbH.

#### 4.7.2 Peptide Preparation

Peptide (ENSFPFFNENSFPFFNENSFPFFN-NH_2_) was synthesized via Fmoc-based SPPS on a CEM Liberty Blue microwave-accelerated peptide synthesizer. Synthesis was performed on a 0.1 mmol scale using TentaGel^®^ S RAM resin (0.23 mmol/g). Five equivalents each of amino acid, Oxyma Pure, and DIC were heated at 90 °C for 4 min to facilitate coupling. Deprotection was achieved using 20% piperidine in DMF heated to 90 °C for 1 min.

Between deprotection and coupling steps, three DMF washes were performed, with a single washing step between the coupling and deprotection steps. The resin was washed three times with DMF, MeOH, and DCM, followed by air drying.

Cleavage was performed with 10 mL TFA containing 2.5% (v/v) Milli-Q water and 2.5% (v/v) TIS for 2 h, followed by precipitation of the product in Et_2_O. The product was collected by centrifugation (4000 rpm, 10 min), the organic layer removed, and the resulting pellet resuspended in 0.1% (v/v) ammonium hydroxide (NH_4_OH) for direct purification.

Peptides were purified using reversed-phase HPLC on a Shimadzu system equipped with two KC-20AR pumps and an SPD-20A or SPD-M20A detector, fitted with a 21.2 × 150 mm Phenomenex Kinetex Evo C18 column. A linear gradient from 10–90% MeCN in water was used at a flow rate of 12 mL/min. Collected fractions were analyzed via analytical HPLC, pooled, and lyophilized twice to yield the dry products. MS characterization of purified peptides can be found in Figure S10.

For peptide sample preparation, peptide was dissolved in 0.1% (v/v) NH_4_OH, and 0.1 M HCl was added to adjust pH to approximately 7.6 (final NH_4_Cl concentration: 20–25 mM). Then, Milli-Q water was added to achieve a final peptide concentration of 0.5 mM. Samples were incubated at 37 °C for various times.

#### 4.7.3 Thioflavin-T (ThT) Fluorescence Assay

ThT fluorescence was measured using a TECAN Spark^®^ microplate reader. ThT was dissolved in Milli-Q water to make 10 mM stock solutions, stored at 4 °C, protected with foil for later use. Before measurement, the 10 mM ThT stock was diluted to 100 µM. Then, 5 µL of the 100 µM ThT solution was mixed with 100 µL peptide sample (0.5 mM) in a flat black 96-well plate. After 10 minutes of incubation, the ThT fluorescence intensity was measured by excitation at 440 nm and emission at 490 nm.

#### 4.7.4 Transmission Electron Microscopy (TEM)

TEM imaging was performed on a JEM1400 Plus (JEOL) transmission electron microscope at 80 kV. Peptide samples were gently mixed with Milli-Q water (1:1, v/v), and 10 µL was placed on Formvar/Carbon-coated copper grids for 90 s, stained with 1 wt% uranyl acetate for 60 s, blotted to remove excess liquid, and air-dried prior to imaging.

#### 4.7.5 Circular Dichroism (CD)

CD spectra were collected on a JASCO J-1500 spectropolarimeter using a 0.1 mm quartz cuvette at 37 °C. Spectra were recorded between 190 nm and 260 nm at 0.2 nm intervals and a 50 nm/min scan rate, with ten subsequent spectra averaged to minimize noise. The CD signal of the NH_4_Cl buffer background was subtracted from sample spectra. The mean residue molar ellipticity *θ* (deg cm^2^ (dmol res)^−1^) was calculated using:

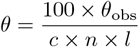

where *θ*_obs_ is the observed ellipticity (mdeg), *c* is peptide concentration (mM), *n* is the number of peptide bonds, and *l* is the path length of the cuvette in cm.

## 5 Acknowledgments

## Author Contributions

A.H., H.J.R., and G.J.A.S. conceived and conceptualized the study. A.H. performed all simulations and computational analyses and prepared the figures. S.Z., E.P., and M.G. performed and analyzed the experiments. A.H., H.J.R., and G.J.A.S. wrote the manuscript. All authors reviewed and approved the final manuscript. H.J.R., G.J.A.S., and A.K. supervised the project.

## 6 Data Availability

The data supporting the findings of this study are available to reviewers as a zipped archive during peer review and will be made publicly available via Zenodo upon publication. All scripts, analysis pipelines, and Evo-MD source code used in this work are available at the following GitHub repository: https://github.com/arthoti/the_missing_link/tree/main.

