## Supplementary Information for "The missing link between biomolecular condensates and amyloid fibrils"

### Supplementary Materials for
### The missing link between biomolecular condensates and amyloid fibrils

Art Hoti*, Shuya Zang, Ephraim Prantl, Michele Grünewald, Alexander Kros,
H. Jelger Risselada*, G. J. Agur Sevink*

**This PDF file includes:**

Supplementary Methods
Figures S1 to S13
Table S1

#### Supplementary Methods

##### Evolutionary Molecular Dynamics (Evo-MD)

Within our Evolutionary Molecular Dynamics (Evo-MD) (*37, 100*) implementation for the design of lyotropic phase forming peptides, coarse-grained molecular dynamics simulations were performed using the Martini 3 force field (*40*) with GROMACS 2023.3 (*103*). Evo-MD was implemented in Python 3.11.3 using MPI4py and NumPy. Coarse-grained peptide models were generated using `martinize2` (*104*) with Martini 3 mapping and no elastic network.

Evo-MD generated an initial population of 512 peptide sequences. Each sequence was constructed by randomly selecting an 8-residue segment and repeating it three times to form a 24-residue peptide composed of three identical repeats. Amino acids were drawn according to probabilities derived from the DisProt intrinsically disordered protein database (*101*). To ensure intrinsic disorder, each sequence was screened using the Porter5 secondary-structure prediction tool (*102*). Sequences for which all residues were annotated as coil were retained, while sequences containing predicted helix or strand were discarded and resampled until a full population of 512 accepted sequences was obtained.

Following initialization, all sequences were subjected in parallel to an automated simulation and analysis pipeline to compute a lyotropic fitness score. The top 64 scoring sequences were selected as parents for recombination. Of these, the top 32 sequences were retained as elites and passed unchanged to the next generation and re-evaluated in the subsequent iteration to reduce stochastic noise in fitness estimation ($N_{\text{parent}} = 64$, $N_{\text{elite}} = 32$). New sequences were generated via uniform crossover between parent sequences, followed by point mutations applied independently to each residue with probability $1/N$, where $N = 24$. Sequences were again filtered through the Porter5 intrinsic disorder filter, until a population of 512 sequences was constructed with the genetic operations. This selection–recombination–mutation cycle was repeated for 20 iterations. Further algorithmic details of the Evo-MD framework are described in previous work (*100, 111*).

Each sequence generated by Evo-MD was processed through four sequential modules: (i) *generate-peptide*, which produced a coarse-grained representation of a single peptide; (ii) *insert-peptide*, which constructed a slab system containing 25 peptide copies; (iii) *production*, which carried out equilibration and production molecular dynamics simulations; and (iv) *compute-fitness*, which analyzed the resulting trajectories to quantify lyotropic ordering.

**Module: generate-peptide** An atomistic template of each *de novo* peptide was generated using PeptideBuilder (*112*) with fixed backbone dihedral angles ($\phi = -60°$, $\psi = -40°$). This atomistic structure served solely as a connectivity template for coarse-graining; subsequent energy minimization and equilibration remove any memory of the initial backbone conformation. The atomistic model was converted to a Martini 3 coarse-grained representation using `martinize2` without an elastic network. The resulting coarse-grained peptide was centered in a $1.65 \times 1.65 \times 20$ nm simulation box, with its long axis aligned along the $z$-axis.

**Module: insert-peptide** Twenty-five individual peptide molecules were arranged into a $5 \times 5$ lattice using the GROMACS `genconf` tool, yielding an initial simulation box of approximately $8.25 \times 8.25 \times 20$ nm. The system was relaxed using steepest-descent energy minimization followed by a short stochastic dynamics simulation (10 ns, 20 fs timestep) to remove steric clashes and relax the imposed lattice geometry. The system was subsequently solvated using the default Martini 3 water model and neutralized with $Na^+$ and $Cl^-$ ions.

**Module: production** Energy minimization was performed using the steepest-descent algorithm for a maximum of 5,000 steps or until the maximum force fell below 10 kJ $mol^{-1}$ $nm^{-1}$. The system was then equilibrated for 250 ps using a stochastic dynamics (Langevin) integrator with a 10 fs timestep. Temperature was maintained at 300 K using the built-in Langevin thermostat with separate coupling groups for protein and solvent ($\tau$ = 1.0 ps). Semi-isotropic pressure coupling was applied using the Berendsen barostat with a reference pressure of 1 bar, a coupling constant of 2.0 ps, and a compressibility of $4.5 \times 10^{-5}$ $bar^{-1}$.

Production simulations were carried out for 400 ns using stochastic dynamics integration with a 20 fs timestep. Temperature and pressure coupling parameters were identical to those used during equilibration. Coordinates were saved every 2 ns for subsequent analysis. Periodic boundary conditions were applied in all three dimensions, and no positional or orientational restraints were used during equilibration or production.

Nonbonded interactions were treated using standard Martini 3 settings. Neighbor searching employed the Verlet cutoff scheme with a cutoff distance of 1.0 nm and neighbor lists updated every 20 steps. Electrostatic interactions were computed using the reaction-field method with a cutoff of 1.1 nm and a relative dielectric constant of 15, while van der Waals interactions were truncated at 1.1 nm. Both potentials were shifted to zero at the cutoff. Bond constraints were enforced using the LINCS algorithm with an expansion order of 8 and two iterations. Center-of-mass motion was removed every 100 steps using linear removal.

**Module: compute-fitness** Number-density profiles of polar and non-polar residues were computed along the $z$-axis using the GROMACS `gmx density` tool with number-density output. Polar residues were defined as ASN, GLU, GLN, THR, SER, CYS, LYS, ARG, HIS, and TYR, while non-polar residues were defined as GLY, ALA, VAL, LEU, ILE, MET, PHE, PRO, and TRP. Density profiles were computed from 100 ns onward, such that the final 300 ns of each trajectory contributed to fitness evaluation. Trajectories were centered on the protein prior to density calculation, and residue selections were defined using index files. Densities were evaluated over the full

$z$-dimension of the simulation box and normalized by the total number density of the corresponding residue class.

The lyotropic fitness was computed as

$$\text{fitness} = -\sum_i \min\left[\frac{\rho_{\text{nonpolar}}(z_i)}{\sum_j \rho_{\text{nonpolar}}(z_j)}, \frac{\rho_{\text{polar}}(z_i)}{\sum_k \rho_{\text{polar}}(z_k)}\right], \tag{S1}$$

where $\rho_{\text{nonpolar}}(z)$ and $\rho_{\text{polar}}(z)$ denote the number densities of non-polar and polar residues, respectively, at position $z$ along the membrane normal.

#### Analysis of solution space and sequence properties

**Population entropy and sequence properties** Sequence entropies (Fig. 1B) were calculated by first computing the Shannon information entropy at each amino acid position across the entire sequence population, then averaging these values across all positions. The Shannon entropy at position $i$ was calculated as $H_i = -\sum_{a=1}^{20} p_{i,a} \log_2 p_{i,a}$, where $p_{i,a}$ is the relative frequency of amino acid $a$ at position $i$. The overall sequence entropy was then computed as the mean of $H_i$ across all positions. SCD (*44*), SHD (*45*), and aromatic clustering (*46*) for each sequence were computed according to the referenced works.

**Statistical Analysis of Compositional Evolution** To quantify evolutionary dynamics and inter-population relationships in amino acid composition, we employed three complementary statistical measures.

Jensen–Shannon (JS) divergence is an entropy-based, information-theoretic metric derived from the Kullback–Leibler (KL) divergence that quantifies distributional differences between amino acid frequency profiles. For two population frequency distributions $P$ and $Q$, JS divergence was defined as

$$\text{JS}(P, Q) = \frac{1}{2}\left[\text{KL}(P\|M) + \text{KL}(Q\|M)\right], \tag{S2}$$

where $M = \frac{1}{2}(P + Q)$ is the average distribution and

$$\text{KL}(P\|Q) = \sum_i P(i) \log_2\left(\frac{P(i)}{Q(i)}\right) \tag{S3}$$

is the standard KL divergence. JS divergence ranges from 0 (identical distributions) to 1 (maximally distinct distributions) and provides a symmetric measure of compositional distance.

Pearson's product–moment correlation coefficient quantified linear relationships between amino acid frequencies across populations:

$$r(P,Q) = \frac{\sum_i \left(P(i) - \bar{P}\right)\left(Q(i) - \bar{Q}\right)}{\sqrt{\sum_i \left(P(i) - \bar{P}\right)^2 \sum_i \left(Q(i) - \bar{Q}\right)^2}}, \tag{S4}$$

where $\bar{P}$ and $\bar{Q}$ denote the mean frequencies. This parametric measure captures proportional covariation in absolute frequency values.

Spearman's rank correlation coefficient assessed monotonic relationships based solely on the ordering of amino acid abundances. For each population, amino acids were assigned ranks from 1 (most abundant) to 20 (least abundant). The rank difference for amino acid $i$ was defined as

$$d_i = r_{P,i} - r_{Q,i},$$

where $r_{P,i}$ and $r_{Q,i}$ denote its rank in distributions $P$ and $Q$, respectively. Spearman correlation was then computed as

$$\rho(P,Q) = 1 - \frac{6 \sum_i d_i^2}{n(n^2 - 1)}, \tag{S5}$$

with $n = 20$ amino acids. This non-parametric measure therefore evaluates similarity in the *ordinal structure* of amino acid composition—that is, the relative ranking of amino acids from most to least frequent—independent of their absolute frequencies.

Together, these three measures describe different ways in which amino-acid compositions can differ between populations. Jensen–Shannon divergence captures differences in the overall shape of the distributions. Pearson correlation measures whether the amino-acid frequencies in one population vary in a proportional, approximately linear way relative to the other. Spearman correlation measures how similarly the amino acids are ranked by abundance, independent of their absolute values.

#### Non-Evo-MD simulations and analysis

##### Large membrane patch

**Simulations** Large-scale membrane systems (32×32 nm, 900 peptide chains) were constructed from equilibrated slab configurations generated during the genetic algorithm optimization. Starting from the final frame of the 5.6×5.6×18 nm slab simulation, solvent molecules were removed and peptide chains were made whole across periodic boundaries using the GROMACS `gmx trjconv` tool. Thirty-six copies of this system were then tiled in a 6×6 array along the $x$ and $y$ dimensions to create the expanded membrane system.

The assembled system was relaxed prior to solvation through energy minimization using the steepest descent algorithm followed by a brief equilibration. The system was then solvated using `gmx solvate` and neutralized with counterions using `gmx genion`.

Production simulations were performed in the $NP_{xy}T$ ensemble with semi-isotropic pressure coupling. Following equilibration (25 ns with a 10 fs timestep), production runs were conducted for 1 $\mu$s using a 20 fs timestep. All other simulation parameters (temperature coupling, electrostatics treatment, force field, and nonbonded interaction settings) were matched to those used in the Evo-MD slab simulations described above, unless stated otherwise.

***Note on large-membrane pressure coupling.*** For the membrane patch simulations of peptide $[ENSFPFFN]_3$, Pressure was controlled using the semi-isotropic Parrinello–Rahman barostat at 1.0 bar with a coupling constant of 12.0 ps and a compressibility of $3 \times 10^{-4}$ bar$^{-1}$. The $[NQSGFFFS]_3$ membrane patch simulation included in the Supplementary Information was generated using an earlier version of the protocol where pressure was controlled using the semi-isotropic Berendsen barostat at 1.0 bar with a coupling constant of 2.0 ps and a compressibility of $4.5 \times 10^{-5}$ bar$^{-1}$ was used. Importantly, the conclusions drawn from these simulations were robust with respect to this protocol variation.

**Structural and dynamical analysis of peptide membranes** All trajectory analyses were performed using the MDAnalysis package (*105, 113*). Trajectories were loaded from GROMACS `.tpr` and `.xtc` files, and backbone atoms were unwrapped using MDAnalysis transformations to remove periodic boundary artifacts.

***Peptide alignment analysis***

The orientation of peptide chains relative to the membrane normal ($z$-axis) was characterized using two complementary metrics. End-to-end vectors were calculated for each 24-residue peptide by selecting backbone beads (`name BB`) and computing the vector between the N-terminal and C-terminal backbone beads. Vectors were normalized by their magnitude, and the angle $\theta$ between each end-to-end vector and the membrane normal was calculated as $\theta = \arccos(\hat{e}_{2e} \cdot \hat{z})$.

In addition, the orientation of phenylalanine aromatic rings was assessed by calculating the normal vector to the ring plane (defined by three sidechain beads) and determining its angle relative to the $z$-axis.

For both metrics, the nematic order parameter was calculated as

$$S = \frac{1}{2}\left(3\langle\cos^2\theta\rangle - 1\right),$$

where $S = 1$ indicates perfect alignment with the membrane normal, $S = 0$ indicates random orientation, and $S = -0.5$ indicates perpendicular orientation. To assess in-plane organization, the $x$–$y$ projections of end-to-end vectors were analyzed for peptides that did not span periodic boundaries, identified by checking whether atomic coordinates spanned more than 50% of the box dimension.

***In-plane nematic order parameter***

For each peptide, the end-to-end vector was projected onto the $x$–$y$ plane and the azimuthal angle $\phi$ was computed as $\phi = \arctan 2(y, x)$. The in-plane nematic order parameter was calculated as

$$S = \left|\left\langle e^{2i\phi}\right\rangle\right|,$$

which yields values between 0 (random orientations) and 1 (perfect in-plane alignment).

***Center-of-mass trajectory analysis***

Center-of-mass trajectories were computed for each peptide by averaging the positions of all backbone beads at each timepoint. Periodic boundary conditions were accounted for by unwrapping trajectories and correcting displacement jumps exceeding half the box length. Only the $x$–$y$ components of the trajectories were retained for in-plane diffusion analysis, and peptides crossing boundaries in the initial frame were excluded.

***Diffusion coefficient calculations***

Two-dimensional translational diffusion coefficients were extracted from mean squared displacement (MSD) analysis of the center-of-mass trajectories, defined as

$$\mathrm{MSD}(\tau) = \langle|r(t+\tau) - r(t)|^2\rangle.$$

Diffusion coefficients were obtained from linear fits to the MSD in the diffusive regime using the Einstein relation MSD = $4D\tau$.

***Backbone dihedral flexibility***

Backbone conformational flexibility was quantified by calculating dihedral angles formed by four consecutive backbone beads along each peptide chain. Circular statistics were used to compute the mean and variance of dihedral angle distributions across sampled frames, with higher circular variance indicating greater flexibility.

***Phenylalanine clustering and hydrophobic core analysis.*** Hydrophobic core formation in lyotropic assemblies was quantified by analyzing collective clustering of phenylalanine (Phe) residues using a connected-component approach. For each trajectory frame, Phe sidechain center-of-mass distances were used to construct a residue–level contact network, and the largest connected component was defined as the hydrophobic core. Core stability and dynamics were quantified from the fraction of Phe residues remaining in the core, Phe–Phe contact statistics, and core membership exchange between consecutive frames.

***Bending rigidity analysis***

Bending rigidities ($\kappa$) were computed from the coarse-grained large membrane patch simulations by analyzing thermal undulations of the membrane midplane. Upper and lower membrane leaflets were identified using leaflet assignments generated with the MDVoxelSegmentation framework (*107*). Instantaneous leaflet height fields were reconstructed from backbone bead positions, and the membrane midplane was defined as the average of the two leaflet surfaces.

Undulation spectra were obtained from the two-dimensional Fourier transform of the midplane height field and radially averaged as a function of wavevector magnitude $q$. Bending rigidities were extracted from the low-$q$ regime assuming a tensionless membrane, using the Helfrich relation $\langle |h(q)|^2 \rangle = k_{\mathrm{B}}T/(\kappa q^4)$. The undulation analysis was performed using an adapted implementation of the method described by Fowler *et al.* (*108*).

#### Solvent-quality scaling simulations

To modulate solvent quality in the coarse-grained simulations, we employed the $\lambda$-scaling protocol of Thomasen *et al.* (*51*), in which all protein–water Lennard–Jones interactions are scaled by a factor $\lambda$. This procedure effectively rescales the depth of the interaction potential between protein and water beads while leaving all other interactions unchanged. By increasing $\lambda$, the Martini beads are made more hyodrophillic.

For each sequence, the starting configuration was obtained from the final frame of the corresponding Evo-MD slab simulation (dimensions ~8×8×20 nm). Solvent molecules were removed, and peptide chains were made whole across periodic boundaries. The resulting assembly was centered in a 25×25×25 nm simulation box, which was chosen to prevent elongated lyotropic structures from interacting with their periodic images. The system was solvated, neutralized, and energy-minimized using the same parameters as in the Evo-MD simulations, followed by a short equilibration run to ensure stability.

At each value of $\lambda$, the system was simulated for 500 ns in the isotropic NPT ensemble using the leap-frog integrator with a 20 fs timestep. Temperature was maintained at 300 K using the velocity-rescaling thermostat with separate coupling for protein and solvent groups ($\tau$ = 1.0 ps). All other simulation parameters – neighbor searching, electrostatics, van der Waals interactions, and constraints – were identical to those used in the Evo-MD slab simulations, with the sole difference being the use of isotropic rather than semi-isotropic pressure coupling. Coordinates were saved every 200 ps for subsequent analysis. Morphologies were classified by visual inspection, and 500 ns was determined to be sufficient for equilibration at each $\lambda$ value.

##### All-atom simulations

Representative coarse-grained assemblies obtained from the solvent-quality scaling simulations were converted to atomistic resolution to validate the stability and structural features of the peptide assemblies. Frames were selected from isotropic simulations at timepoints where the assembled structures did not cross periodic boundaries. Selected frames were centered in a 17.5×17.5×17.5 nm simulation box to prevent self-interactions across periodic boundaries.

Backmapping was performed using the `backward.py` script with the `initram.sh` wrapper. An atomistic template of a single peptide was first generated using `gmx pdb2gmx` with the a99SB-disp force field, ignoring hydrogens in the input structure. The topology was modified to reflect the appropriate number of peptide chains, and the coarse-grained configuration was then backmapped to atomistic resolution using the Amber99SB mapping scheme.

The backmapped system was solvated using the a99SB-disp water model and neutralized with sodium and chloride ions. Energy minimization was performed using the steepest descent algorithm for up to 50,000 steps with a force tolerance of 1000 kJ $mol^{-1}$ $nm^{-1}$ and an initial step size of 0.01 nm. Electrostatic interactions were treated using the Particle Mesh Ewald (PME) method with a real-space cutoff of 1.0 nm, and van der Waals interactions were truncated at 1.0 nm using the Verlet cutoff scheme.

Following energy minimization, the system was equilibrated for 100 ps in the NVT ensemble using the leap-frog integrator with a 2 fs timestep and position restraints applied to protein atoms. Temperature was maintained at 300 K using the velocity-rescaling thermostat with separate coupling for protein and non-protein groups ($\tau_t = 0.1$ ps). The system was subsequently equilibrated for 100 ps in the NPT ensemble with position restraints maintained, using isotropic Berendsen pressure coupling at 1 bar with a time constant of 2.0 ps.

Production simulations were performed for 400 ns in the NPT ensemble using the leap-frog integrator with a 2 fs timestep. Position restraints were removed for production runs. Temperature was maintained at 300 K using the velocity-rescaling thermostat, and pressure was controlled using the Parrinello–Rahman barostat with a time constant of 2.0 ps and a compressibility of $4.5 \times 10^{-5}$ $bar^{-1}$. Coordinates were saved every 80 ps and energies every 50 ps.

For all atomistic simulations, periodic boundary conditions were applied in all three dimensions. PME electrostatics were used with fourth-order interpolation and a Fourier grid spacing of 0.16 nm. All bonds involving hydrogen atoms were constrained using the LINCS algorithm.

**Ramachandran analysis**

Backbone conformational preferences were assessed through Ramachandran analysis of the $\phi$ and $\psi$ dihedral angles. Dihedral angles were computed for all residues using the MDAnalysis Ramachandran module, with trajectories subsampled every 10 frames. Terminal residues lacking defined dihedrals were excluded. The resulting $\phi$–$\psi$ distributions were analyzed across the full angular range from $-180°$ to $180°$.

**Phenylalanine–phenylalanine contact analysis**

Aromatic interactions between phenylalanine residues were analyzed using MDAnalysis. For each trajectory frame, centroids of phenylalanine aromatic rings (atoms CG, CD1, CD2, CE1, CE2, CZ) were calculated. Pairwise centroid-to-centroid distances were computed with periodic boundary conditions applied. Unique residue pairs were identified using global residue indices to avoid duplication across peptide chains. A distance cutoff of 7 Å was used to define a contact, consistent with established favorable phenylalanine stacking distances (*106*). Contact frequencies were calculated as the fraction of frames in which a given pair satisfied this criterion.

**Hydrogen bond analysis**

Hydrogen bonding patterns were analyzed using the MDAnalysis `HydrogenBondAnalysis` module. Hydrogen bonds were identified using a donor–acceptor distance cutoff of 3.0 Å and a donor–hydrogen–acceptor angle cutoff of 150°. Donors included backbone amide nitrogens and polar sidechain atoms, while acceptors included backbone carbonyl oxygens and polar sidechain heteroatoms. Intra-residue hydrogen bonds were excluded. Trajectories were subsampled every 20 frames, and hydrogen bonds were deduplicated by unique donor–acceptor atom pairs prior to analysis.

### Supplementary Figures

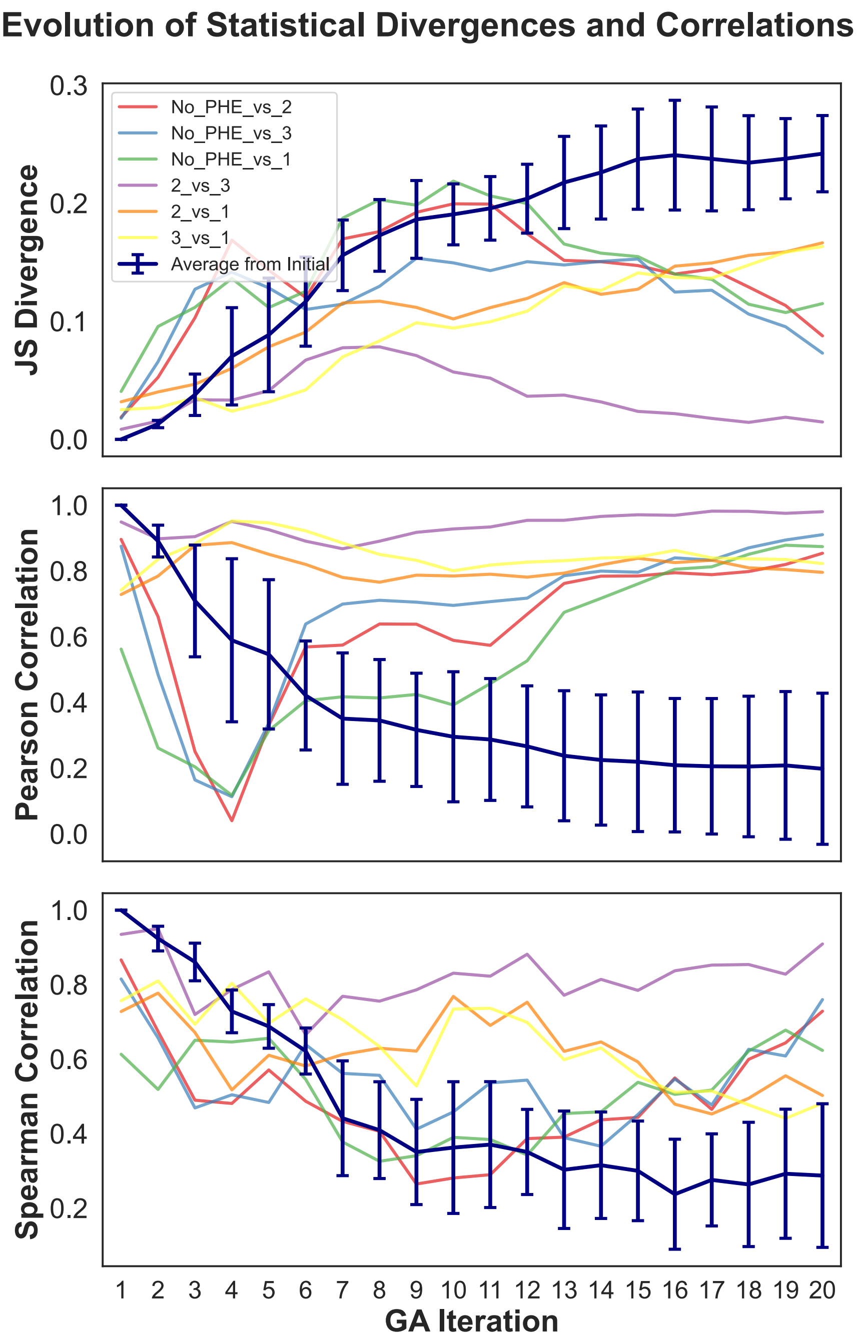


**Figure S1.** Statistical comparison of amino acid frequency distributions for the top 64 scoring peptides at each iteration along each evolutionary trajectory. The solid blue lines show the per-iteration mean across replicas for each statistic, computed between each replica and its own initial state; error bars are ± 1 sample SD across replicas. This initial-state comparison serves as a baseline, so the inter-replica curves can be interpreted in light of whether replicas are drifting similarly or diverging strongly from one another. Colored lines represent pairwise comparisons between replicas at the same iteration; color assignments are indicated in the figure legend. Panels report Jensen–Shannon divergence, Spearman, and Pearson correlations.

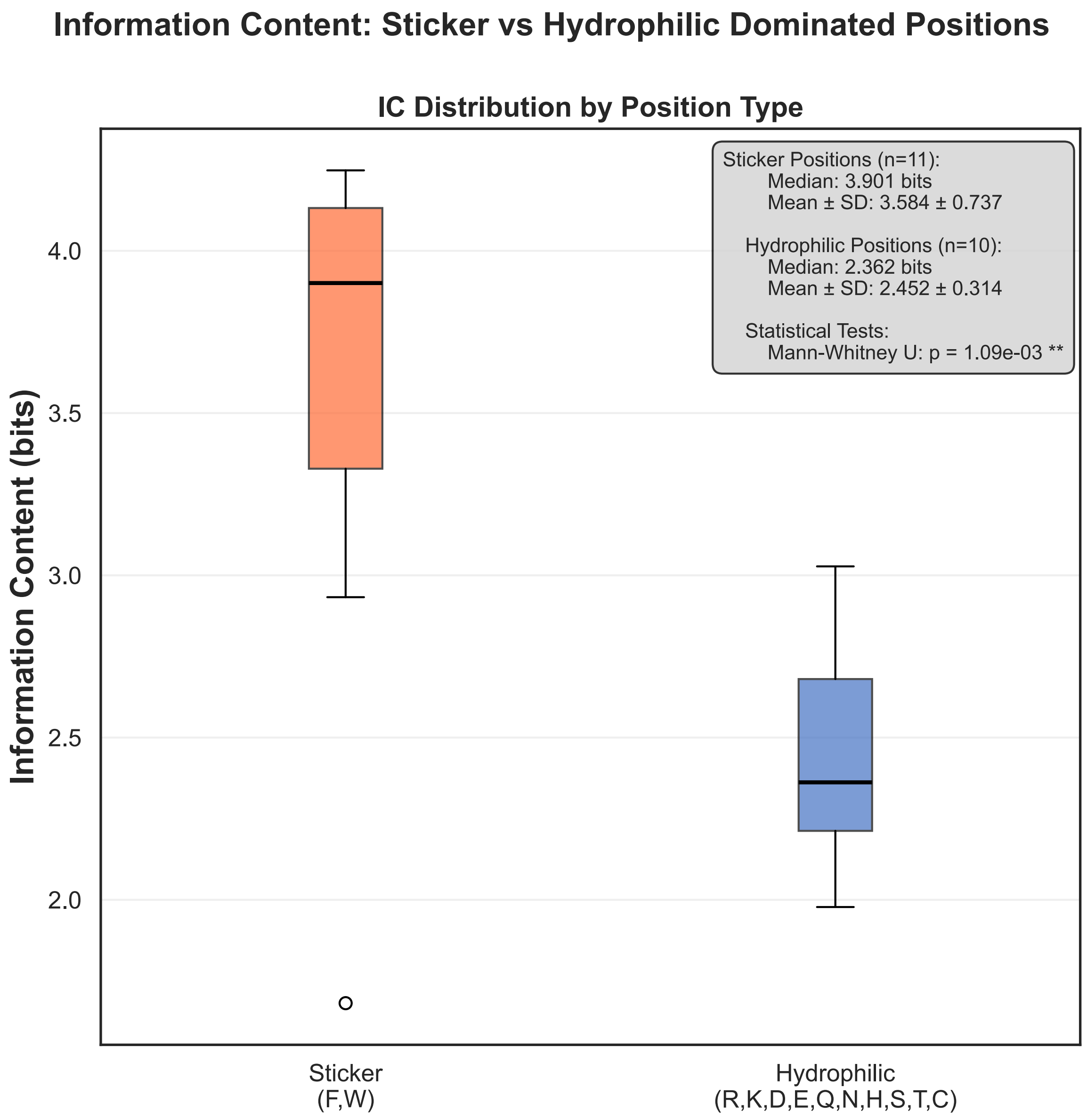


**Figure S2.** Boxplots show positional information content (bits) for sticker-dominated (F/W) and hydrophilic-dominated (R,K,D,E,Q,N,H,S,T,C) sequence positions across replicas. Boxes indicate median and inter-quartile range (IQR); whiskers show 1.5× IQR; outliers are plotted as points. Colors denote groups; the inset reports summary statistics and the Mann–Whitney U test p value.

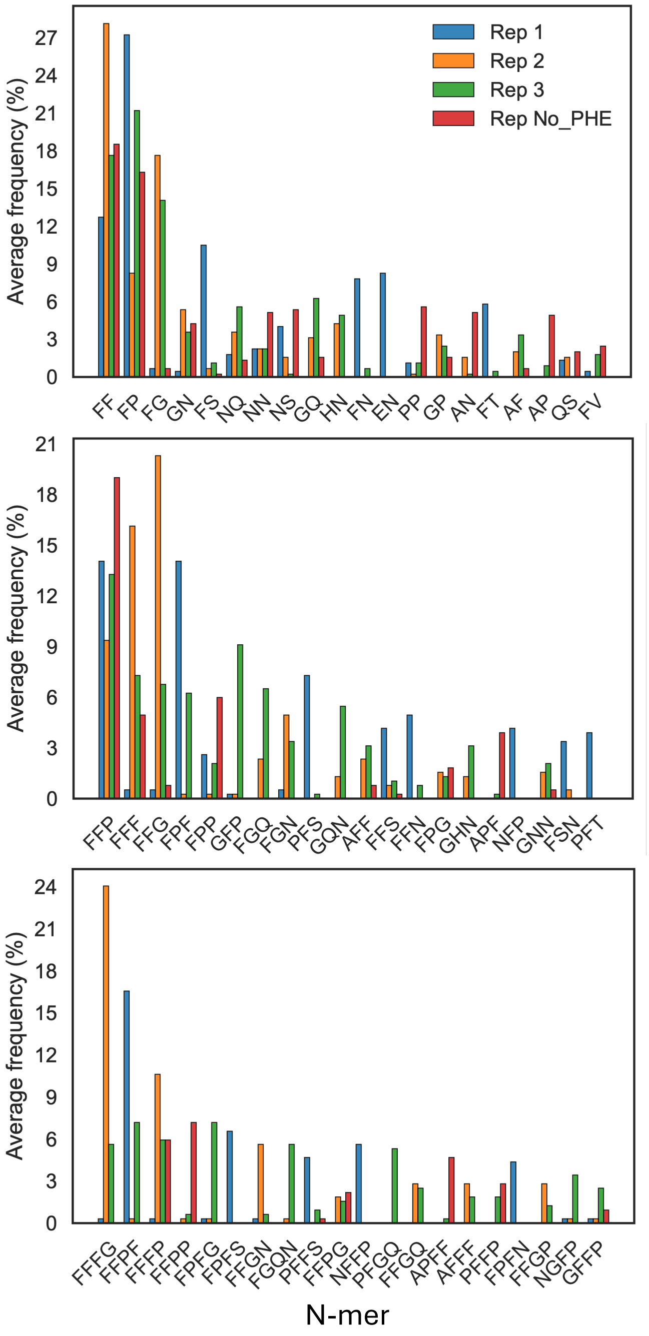


**Figure S3. Frequency distributions of enriched N-mers.** Frequency distributions of the top 20 N-mers among the top 64 scoring sequences in the final iteration of each replica. 2-, 3-, and 4-mers are shown from top to bottom.

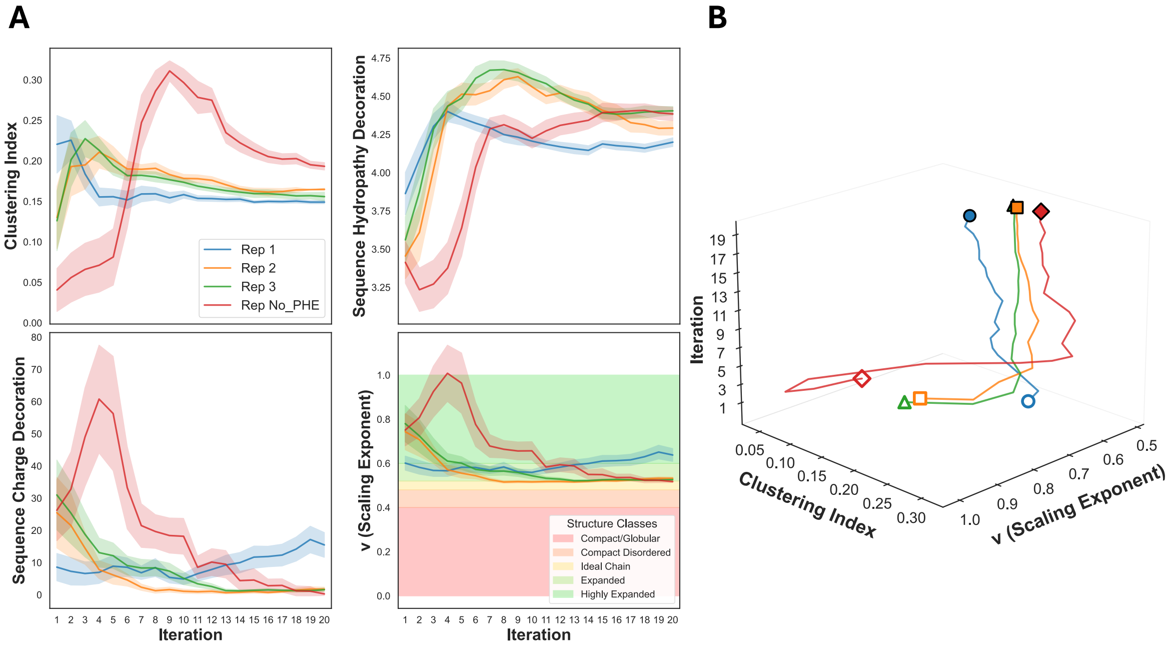


**Figure S4.** (A) Evolution of IDP sequence metrics over 20 iterations of EvoMD. For each iteration, the mean and standard deviation are shown for the top 64 sequences. The scaling exponent plot highlights characteristic chain behavior across different regions of $\nu$ space. (B) Three-dimensional representation of the coevolution of the aromatic clustering index and the scaling exponent $\nu$ across all replicas during the EvoMD optimization procedure.

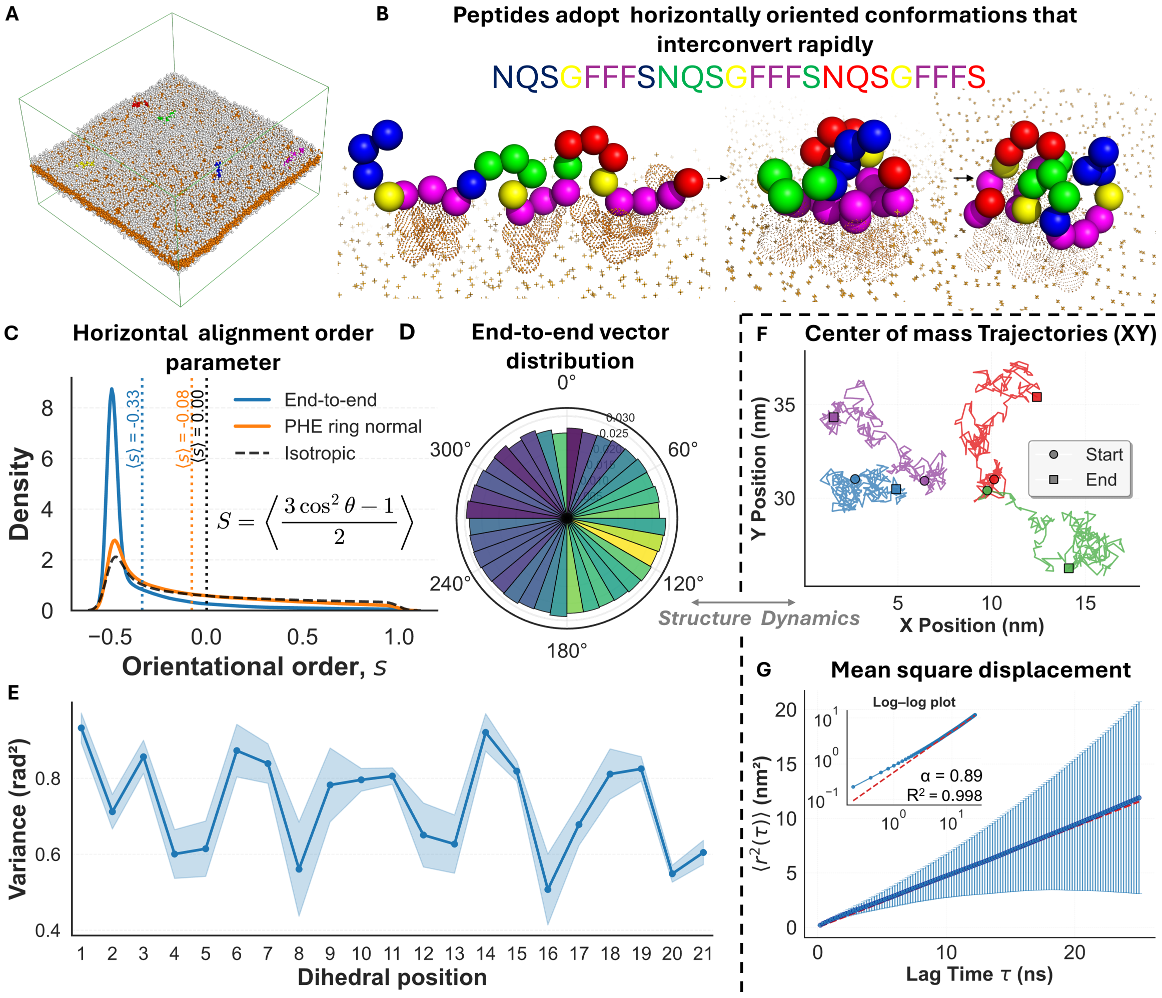


**Figure S5. Structural and dynamic analysis of the $[NQSGFFFS]_3$ membrane system.** (A) Bead representation of a simulated 32×32 nm membrane formed by 900 copies of $[NQSGFFFS]_3$. Hydrophobic residues (orange) and hydrophilic residues (white). Five chains shown in different colors highlight horizontal embedding within the membrane. (B) Three conformations of the same membrane peptide at three different instances along the $1\mu s$ simulation trajectory. Backbone beads are colored according to the sequence displayed above; aromatic Phe sidechains are shown as orange beads. The chain adopts a horizontal conformation with phenylalanine side chains inserted into the hydrophobic core (orange crosses) while the backbone explores different conformations, from extended (left) to compact (right). (C) Distributions of the chain-wise orientational order parameter, $s$. The ensemble average $\langle s \rangle = S$ measures peptide alignment with respect to the membrane normal across all chains and simulation frames. Vertical dotted lines indicate the values of $S$ for $\theta_{\mathrm{E2E}}$,

$\theta_{\mathrm{Phe}}$, and an isotropic distribution, while solid lines show the corresponding $s$ distributions. (D) Rose diagram of peptide chain end-to-end vectors projected onto the $x$–$y$ plane. The angular distribution is approximately isotropic, indicating no preferred alignment. The radial extent of each bin corresponds to the probability that a chain within the ensemble is oriented along the angular range of that bin. The color of each sector encodes the mean end-to-end distance of chains in the bin, ranging from 18.6 Å (dark blue) to 19.8 Å (yellow). (E) Variance of dihedral angles along the peptide backbone reveals a periodic pattern where angles between repeat segments are most constrained (lowest variance). (F) Three representative peptide center-of-mass trajectories in the x-y plane. (G) Mean square displacement plot with log-log inset reveals normal fluid behavior with weak subdiffusion ($\alpha$ = 0.89) in two dimensions.

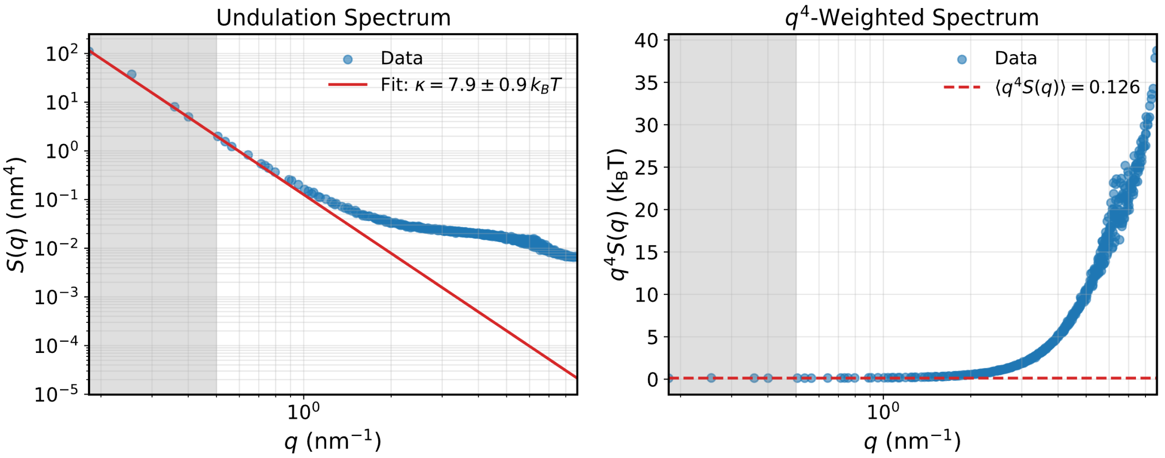


**Figure S6. Determination of membrane bending rigidity.** Undulation spectrum (left) and corresponding $q^4$-weighted spectrum (right) for a membrane formed by peptide $[\mathrm{ENSFPFFN}]_3$. Symbols denote simulation data. Solid lines in the undulation spectra show fits to the Helfrich elastic model in the bending-dominated regime, $\langle |h(q)|^2 \rangle = k_\mathrm{B}T/(\kappa q^4)$. Shaded regions indicate the wavevector range used for fitting. In the $q^4 S(q)$ representation, a plateau identifies the bending regime, with the dashed line indicating the mean value $k_\mathrm{B}T/\kappa$ from which the bending rigidity $\kappa$ is obtained.

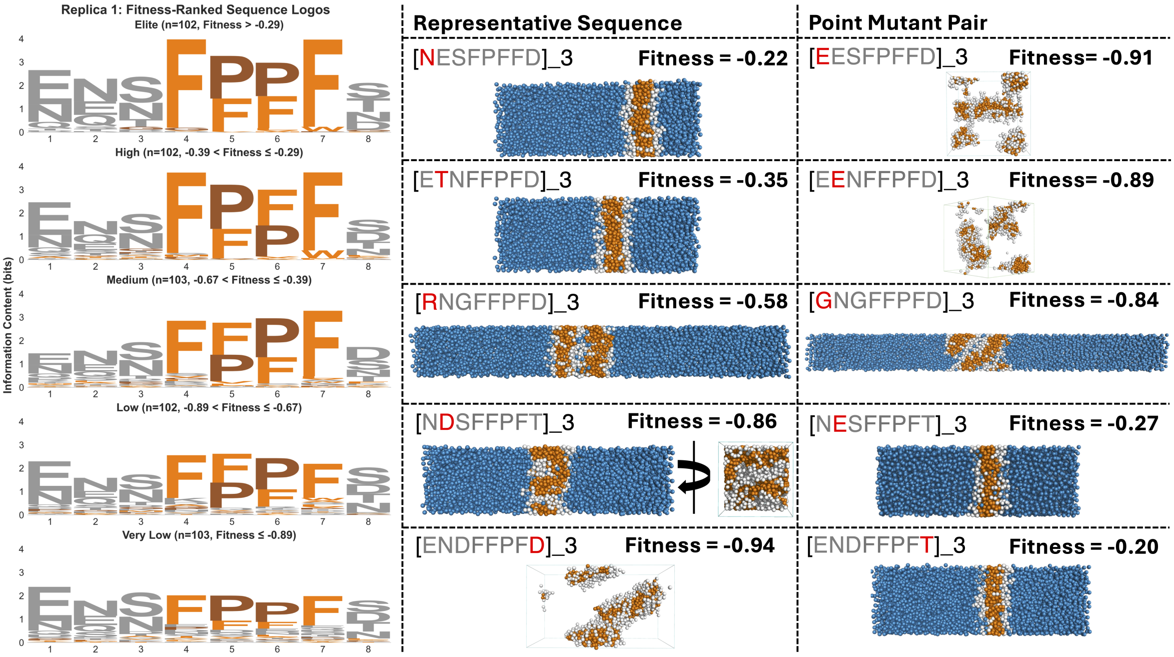


**Figure S7.** (Left) Sequence logos for all 512 sequences from the final iteration of replica 1, grouped into fitness quintiles. Although overall amino acid enrichment is largely conserved across quintiles, the information content at key hydrophobic positions decreases with decreasing fitness. (Right) Point-mutation experiments illustrating the effects of single-residue substitutions on lyotropic morphology and associated fitness in slab simulations. Physicochemical and geometric perturbations introduced by point mutations shift sequences within the morphological phase diagram, producing deviations from lamellar geometries without eliminating lyotropic phase formation. Hydrophilic residue beads are shown in white and hydrophobic beads in orange. For clarity, water beads are displayed in some systems but omitted in others. Accordingly, as evolutionary optimization proceeds and genetic diversity decreases, the population average shifts toward increasingly lyotropic behavior; however, strict lamellar geometries emerge only among the highest-fitness sequences.

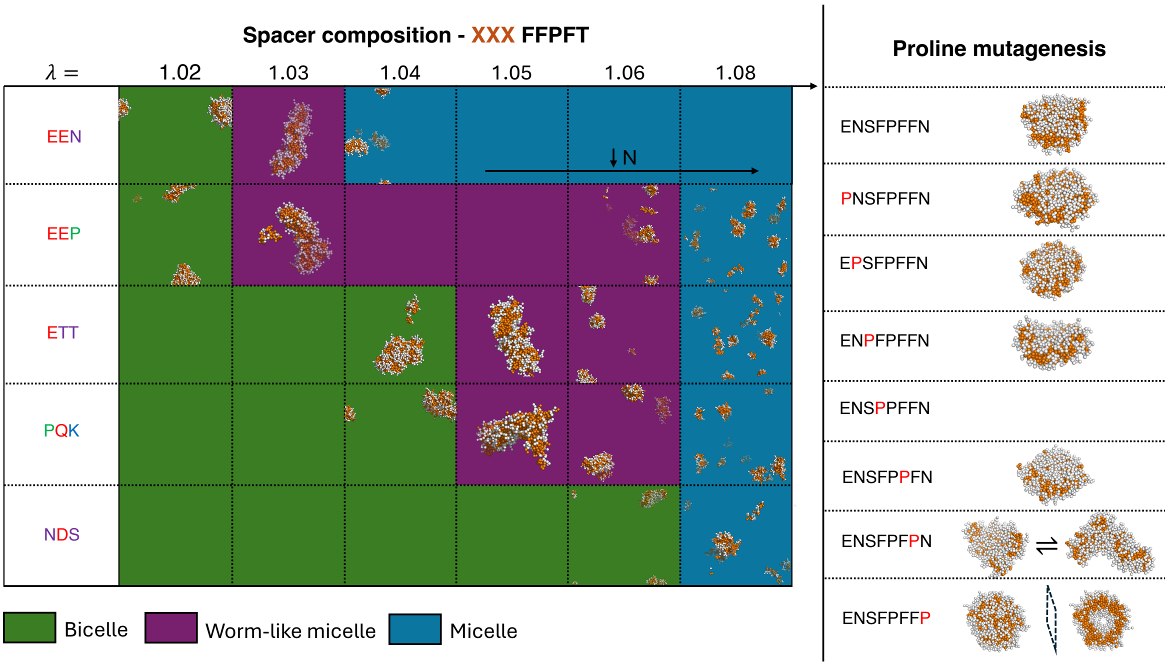


**Figure S8.** (Left) Morphological phase boundaries for sequences of the form [*XXX*FFPFT]$_3$ with varying spacer compositions. Morphologies are mapped as a function of the solvent quality, $\lambda$, shown on the horizontal axis. Altering the identity of the hydrophilic spacer preserves overall lyotropic character but shifts the relative boundaries between morphological regimes. Colors in the legend denote morphology, assigned by visual inspection. Insets show bead representations of the final frames from 500 ns isotropic simulations; to minimize clutter, snapshots are provided only for the first and final $\lambda$ values corresponding to each morphology. Orange beads represent hydrophobic phenylalanine residues, while white beads represent all other residues. (Right) Proline mutagenesis of the elite sequence [ENSFPFFN]$_3$, probing the role of proline-induced torsional constraints. Notably, the sequence [ENSFPFF**P**]$_3$ adopts the appropriate packing geometry to form a hollow vesicle.

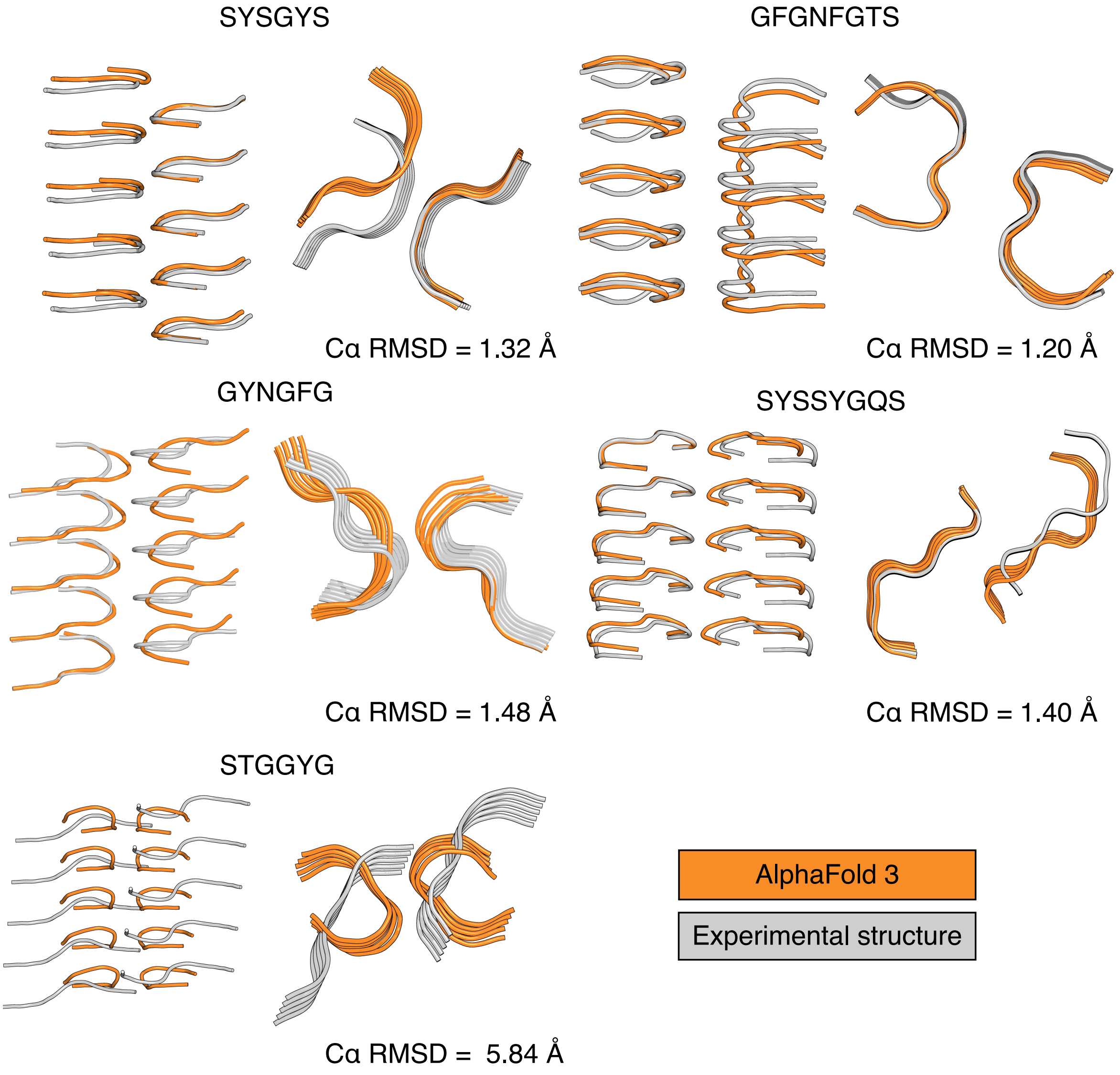


**Figure S9. AlphaFold 3 LARKS benchmark.** Cartoon representations of AlphaFold 3–predicted peptide structures (orange) are aligned to experimentally determined PDB structures (light orange). The benchmark includes the LARKS peptides SYSGYS (PDB ID: 6BWZ), GFGNFGTS (PDB ID: 6BZM), GYNGFG (PDB ID: 6BXX), SYSSYGQS (PDB ID: 6BXV), and STGGYG (PDB ID: 6BZP). Whole-structure C$\alpha$ RMSD values are reported for each alignment.

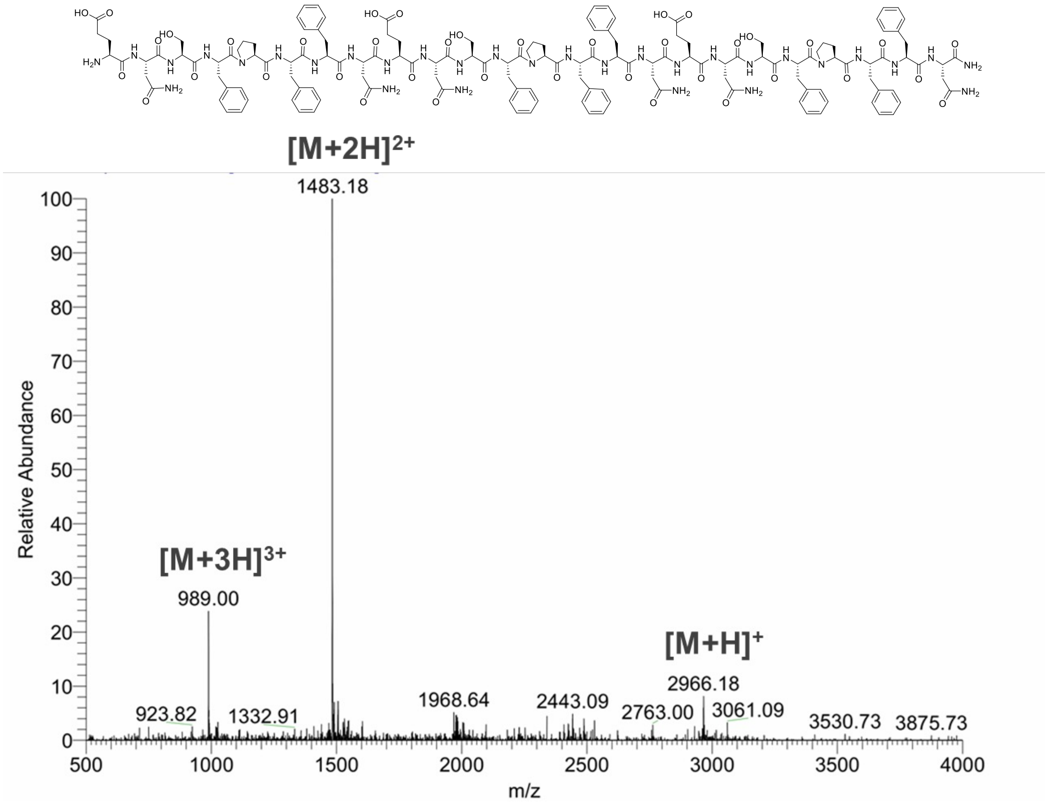


**Figure S10.** (Top) Structural representation of the synthesized [ENSFPFFN]$_3$-$NH_2$ peptide used in the experiments. (Bottom) LC-MS spectrum of the [ENSFPFFN]$_3$-$NH_2$ peptide.

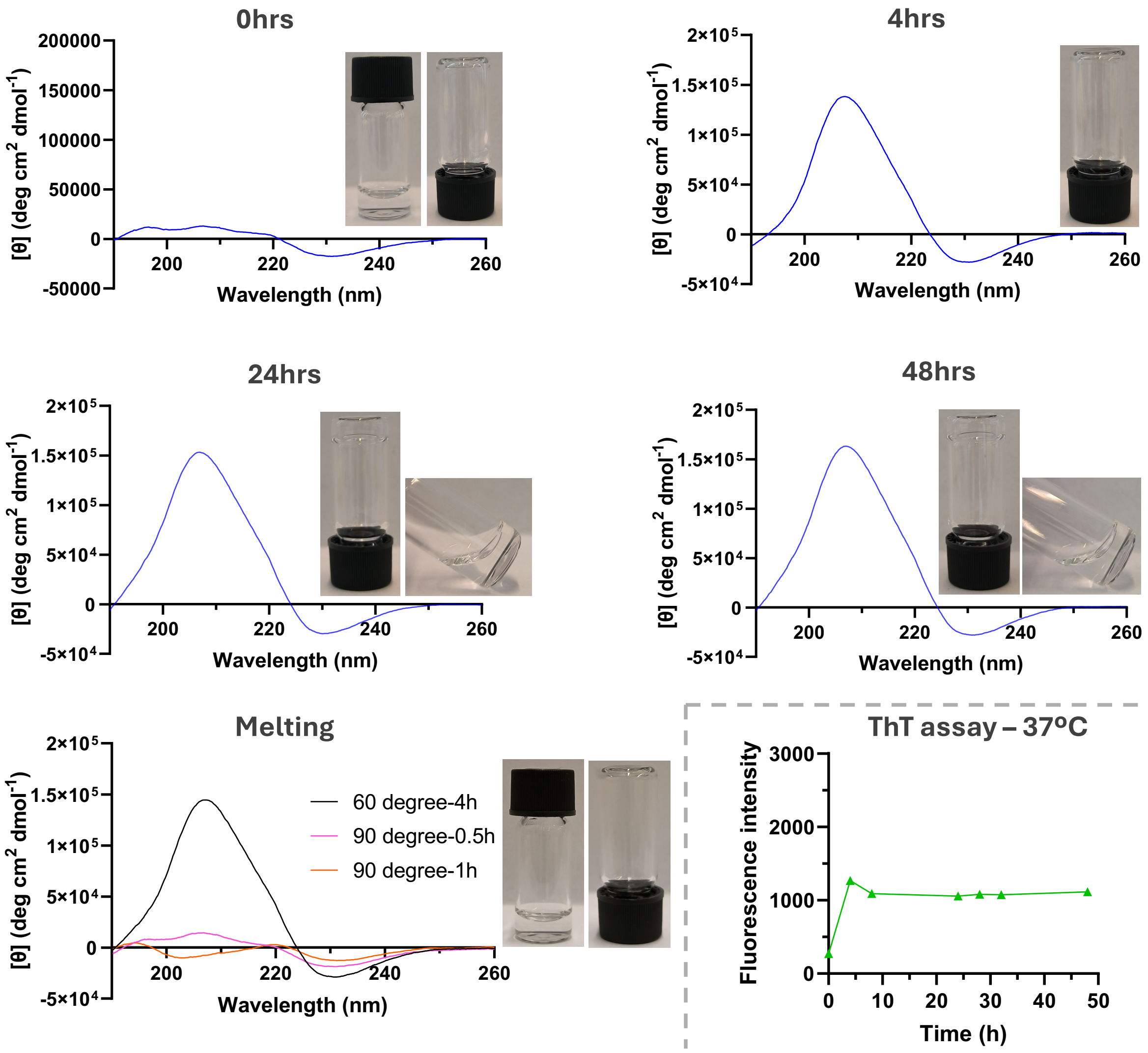


**Figure S11.** Time series of circular dichroism (CD) and inverted vial assays for [ENSFPFFN]$_3$ at 0.5 mM and 37 °C. As the $\beta$-sheet CD signal increases in magnitude, hydrogel formation is observed. Thermal melting experiments demonstrate gel dissolution upon heating to 90 °C, accompanied by a reduction in $\beta$-sheet signal intensity. (Bottom right) Thioflavin T (ThT) fluorescence exhibits a significant signal, consistent with the presence of $\beta$-rich amyloid fibrillar structures observed in TEM and CD measurements.

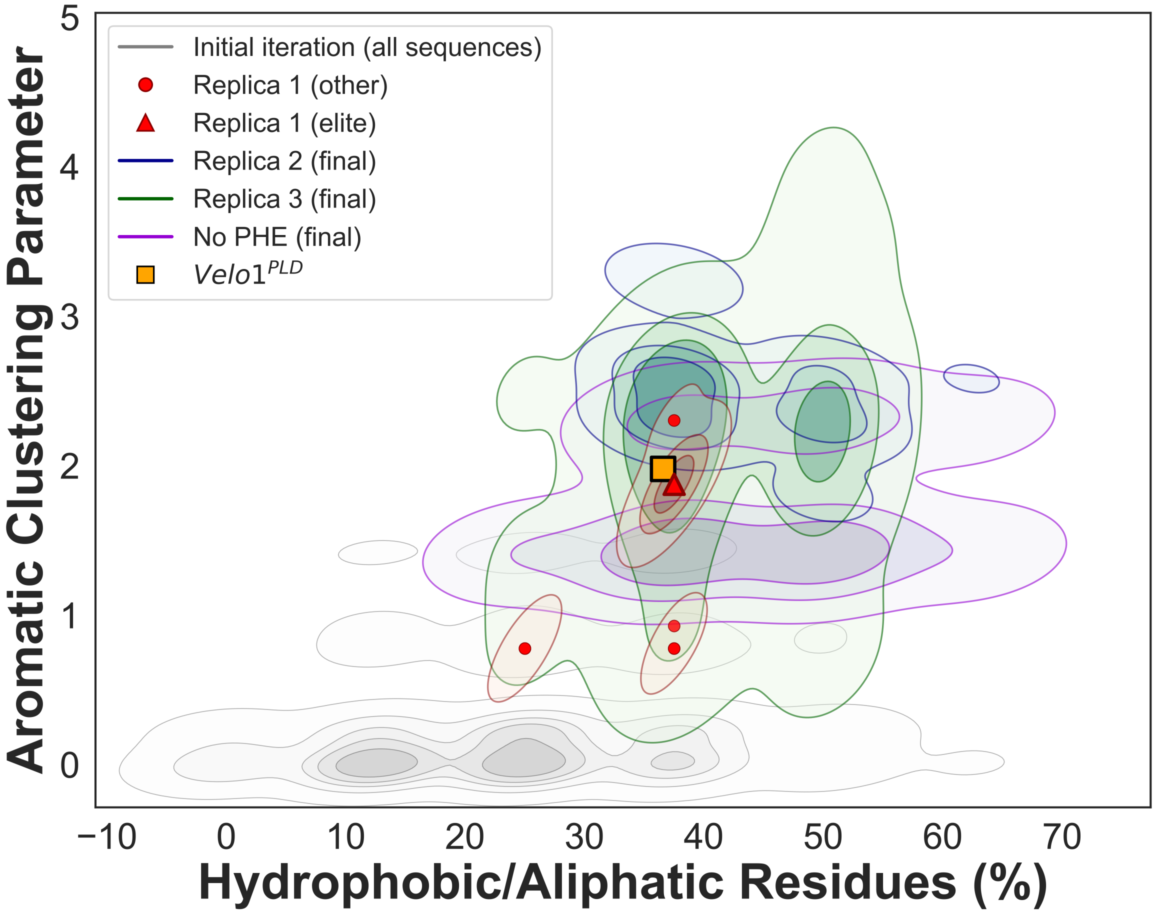


**Figure S12.** Plot illustrating the relationship between aromatic clustering and the fraction of hydrophobic and aliphatic residues in the sequence. The distribution evolves from the lower left (low aromatic clustering and low hydrophobic content) toward the upper right over the course of optimization. The elite sequence is compared to the Balbiani body–forming IDP Velo1$^{PLD}$, which is known to form a functional amyloid-containing condensate. This comparison is inspired by Holehouse *et al.*, who demonstrated a close relationship between these parameters and the material state of the resulting assembly (*46*).

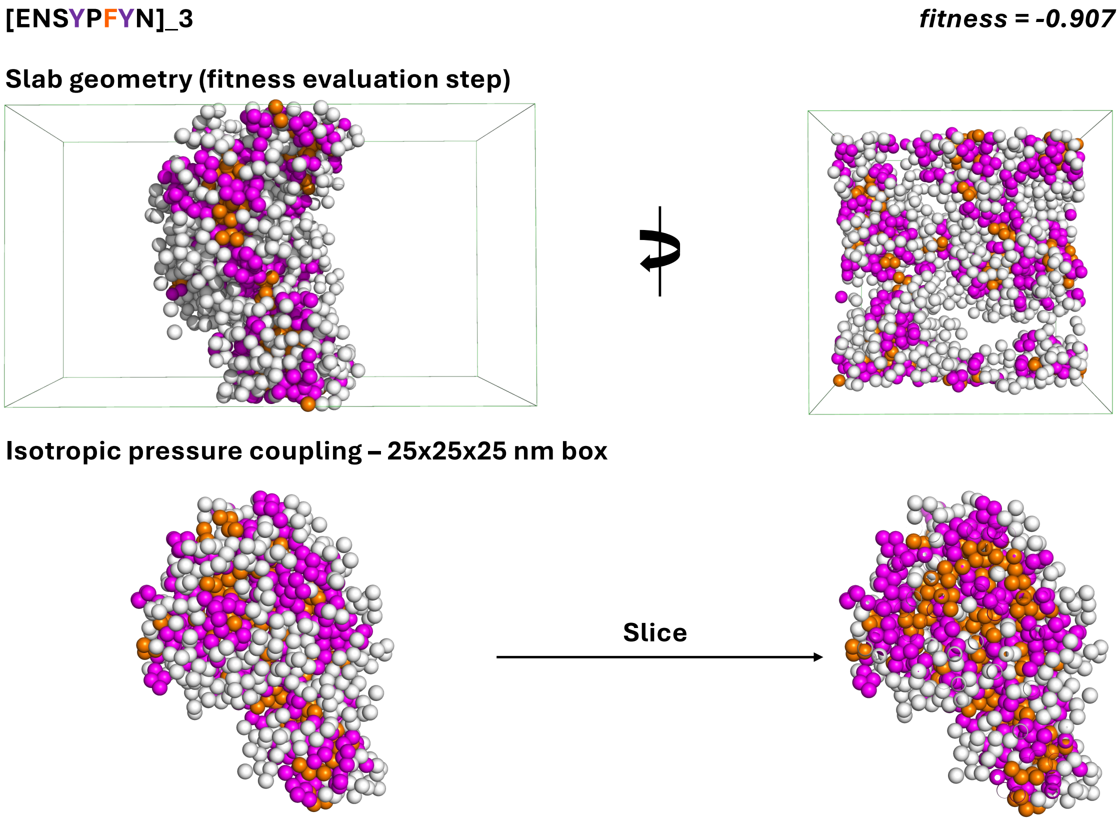


**Figure S13.** Self-assembly of [ENS**Y**PF**Y**N]$_3$ illustrating the effect of tyrosine residues on lyotropic phase morphology. A lyotropic phase forms with tyrosines contributing to the hydrophobic core; however, the phenolic hydroxyl group introduces additional polarity, resulting in a more irregular (jagged) geometry compared to the phenylalanine-only case. Tyrosine residues are shown in magenta, phenylalanine in orange, and all other residues in white.

**Table S1. AlphaFold 3 prediction data for steric zipper, LARKS, and designed peptides.**

Monomer and whole-structure RMSDs, and confidence scores are reported. RMSD values are given for both all-atom and backbone C$\alpha$ atoms where experimental structures are available.

| Sequence | PDB ID | Monomer RMSD | Monomer C$\alpha$ RMSD | Whole RMSD | Whole C$\alpha$ RMSD | Mean pLDDT | iPTM | PTM |
|---|---|---|---|---|---|---|---|---|
| **Steric zipper** | | | | | | | | |
| KLVFFA | 3OW9 | 1.709 | 0.362 | 1.457 | 0.441 | 57.51 | 0.24 | 0.29 |
| **LARKS** | | | | | | | | |
| SYSGYS | 6BWZ | 0.494 | 0.241 | 1.470 | 1.319 | 59.61 | 0.31 | 0.36 |
| GFGNFGTS | 6BZM | 0.756 | 0.250 | 1.852 | 1.200 | 76.15 | 0.64 | 0.66 |
| SYSSYGQS | 6BXV | 0.314 | 0.209 | 1.446 | 1.399 | 62.44 | 0.41 | 0.45 |
| GYNGFG | 6BXX | 1.202 | 0.399 | 3.115 | 1.478 | 73.18 | 0.57 | 0.60 |
| STGGYG | 6BZP | 4.480 | 4.027 | 6.075 | 5.835 | 64.29 | 0.33 | 0.38 |
| **Designed sequences** | | | | | | | | |
| IWWPANNG | – | – | – | – | – | 62.71 | 0.50 | 0.53 |
| ENSFPFFN | – | – | – | – | – | 66.81 | 0.59 | 0.62 |
| NQNGFPFD | – | – | – | – | – | 77.24 | 0.53 | 0.56 |
| ENGFPFFN | – | – | – | – | – | 65.89 | 0.41 | 0.46 |
